# Drainage water management reduces nutrient, *Escherichia coli*, and antibiotic resistance gene export

**DOI:** 10.64898/2026.09.28.755045

**Authors:** Emma M. Thrift-Cahall, Jennifer L. Tank, Ursula H. Mahl, Mitchell J. Liddick, Andrei Badilla-Aguilar, Kyle Bibby

## Abstract

Subsurface tile drainage rapidly transports nutrients and biological contaminants to surface waters, yet multi-year evaluations of drainage water management (DWM) effects on nutrients, *Escherichia coli*, and antibiotic resistance genes (ARGs) remain limited. A paired-field design with high-frequency soil moisture monitoring was used to quantify water yield, dissolved nitrogen, phosphorus, and *Escherichia coli* (*E. coli*) over seven and eight water years, and ARGs (*tetM*, *tetW*, *ermB*, *sul1*) over two years in southern Michigan. Soil moisture at tile depth (∼75 cm) was higher under controlled drainage, confirming that DWM increases field water-holding capacity. Compared to the free drainage field, DWM reduced cumulative tile discharge by 43% and annual nutrient yields by 46-83% for nitrate and 12-82% for soluble reactive phosphorus, but effects on ammonium were variable. Average daily nutrient yields declined by ∼50% over the study period, driven primarily by managed periods. Nutrient concentrations did not differ between treatments, indicating that yield reductions resulted from decreased water loss. Mean daily *E. coli* yields declined by 71% over the study period, and concentrations were 46% lower under controlled drainage. Controlled drainage reduced *tetM* and *sul1* ARG concentrations and yields during managed periods and throughout the study; however, sensitivity analyses showed that these effects were strongly influenced by a high-precipitation event. After excluding the event, differences in ARG concentrations were no longer detected, although ARG yields remained 43% lower during managed periods. These findings highlight the potential for DWM to mitigate nutrient, microbial, and antibiotic resistance transport from agricultural drainage systems.

## 1.0 Introduction

Subsurface tile drainage is implemented on ∼19 million ha of cropland in the United States (USDA NASS, 2022), with 84% of tile-drained croplands concentrated in six Midwestern states representing the “corn belt” (Valayamkunnath et al., 2020). Tile drainage lowers the water table and remove excess soil moisture, improving soil conditions and crop yields (Fausey et al., 1995; Williams et al., 2015), but also accelerates transport of dissolved nutrients and manure-derived microbial contaminants from agricultural fields to adjacent waterways (i.e., streams and ditches; Gaynor et al., 2002; Ross et al., 2016; Skaggs et al., 1982). Producers apply animal manure as a soil amendment and synthetic fertilizers to maximize crop yields, but precipitation (e.g., spring snowmelt and storms) enhances losses of dissolved nutrients (e.g., nitrate; NO_3_^−^-N, ammonium; NH_4_^+^-N, and soluble reactive phosphorus; SRP) and manure-derived pathogens (e.g., *Escherichia coli*, *Salmonella* spp., *Campylobacter* spp.) from tiles. These losses contribute to water quality degradation and eutrophication, with cascading effects on ecosystems, recreation, and livelihoods (Carpenter et al., 1998; Smith et al., 1999; Wilkes et al., 2014). Although tile drainage effects on nutrients and manure-derived microbial residuals have been studied extensively individually (Blann et al., 2009; Frey et al., 2013; Gottschall et al., 2010; Joy et al., 2013; King et al., 2015) these contaminants have rarely been examined in tandem, limiting understanding of how hydrologic management shapes coupled dissolved nutrient and microbial export.

Drainage water management (DWM) uses manually-adjustable weirs to control tile outflow, retaining moisture and nutrients during the fallow period, while allowing free drainage during the growing season and potentially improving crop yields (Ghane et al., 2012; Singh et al., 2022). The effectiveness of DWM on tile flow and nutrient reduction varies with system design, soil type, hydrology, climate, and site conditions (Frankenberger et al., 2023; Skaggs et al., 2012; Williams et al., 2015). Most field evaluations have been conducted over relatively short periods, limiting assessment of interannual variability in weather, hydrology, crop rotations, and manure management that characterize working agricultural landscapes. In a review of 31 studies evaluating DWM effects on nitrogen (N) and/or phosphorus (P) export, the mean study duration was only 3.8 ± 3.2 years (Frankenberger et al., 2023). Consequently, the long-term effectiveness of DWM under the range of conditions encountered on working farms remains understudied. Many DWM studies have focused on NO_3_^−^-N yields, because this form of dissolved nitrogen is highly mobile in water and is a primary driver of downstream eutrophication, as highlighted in several reviews (David et al., 2010; Skaggs et al., 2012). Reductions in NO_3_ ^−^-N yields under DWM are understood to be driven by reduced tile flow and enhanced field-scale denitrification under saturated soil conditions (Review by Frankenberger et al., 2023). In contrast to NO_3_^−^-N, P losses have been less studied as they have historically been assumed to occur mainly through surface runoff and erosion (Dodds and Welch, 2000; King et al., 2015). However, SRP losses from tile drains can be significant, with inconsistent effects of DWM reported across studies (Frankenberger et al., 2023; Osterholz et al., 2024). Though some studies have evaluated NH _4_^+^-N losses (Wang et al., 2020; Sunohara et al., 2015), they have received considerably less attention compared to NO_3_^−^-N and SRP because NH_4_^+^-N typically represents a small fraction of N in tile drainage due to strong soil adsorption and rapid nitrification (Vincent et al., 2025).

In addition to nutrients, manure amendments can introduce bacteria (e.g., *Escherichia coli*; *E. coli*), antibiotic residues, antibiotic resistant bacteria (ARB), and antibiotic resistance genes (ARGs) to agricultural fields, all of which can be transported to nearby waterways via tile drainage (Gottschall et al., 2010; Massé et al., 2014; Bolster et al., 2018; Zhang et al., 2020).

Agricultural land is an important driver of ARG abundance in surface waters, and aquatic ecosystems are increasingly recognized as reservoirs of antibiotic resistance (Allen et al., 2010; Balcázar et al., 2015). Without intervention, deaths from antibiotic-resistant infections are projected to surpass those from HIV, tuberculosis, and malaria combined within 25 years (Naghavi et al., 2024). Although designed primarily to manage water flow and nutrient transport, DWM was not intended to control biological contaminants, which have fundamentally different attributes than solutes; *E. coli* is a bacterial organism capable of growth and replication, whereas ARGs may be carried within bacterial cells or occur as extracellular DNA (Zarei-Baygi and Smith, 2021). By altering subsurface flow, soil saturation, and water residence times, DWM may indirectly influence the persistence and transport of *E. coli*, ARB, and ARGs from agricultural fields (Frey et al., 2013). For example, the persistence of ARGs vary across different soils (Seyoum et al., 2021) and soil conditions like pH, salinity, nutrient content, organic matter, and heavy metals can influence the variation and distribution of ARGs (Wu et al., 2023; Xu et al., 2024). Prior studies have shown that DWM can lower *E. coli* yields through reduced tile flow, although concentration results varied at larger watershed scales (Frey et al., 2013; Schmidt et al., 2013; Sunohara et al., 2016). Critically, only one study has assessed DWM effects on ARGs, representing an ongoing knowledge gap given the growing recognition of agricultural ditches and streams as conduits for antibiotic resistance (Frey et al., 2015). Furthermore, Frey et al. (2015) found that bacterial concentrations do not always mirror nutrient concentrations trends under DWM, suggesting that biological and biogeochemical contaminants may respond differently to drainage management.

To understand the drivers behind dissolved nutrient, *E. coli*, and ARG dynamics, we conducted a multi-year study using a paired field design comparing one field managed with DWM (i.e., controlled drainage; CD) to one with free drainage (FD). We monitored NO_3_^−^-N, NH_4_^+^-N, and SRP over seven water years (WY 2019-2025), *E. coli* and discharge over eight water years (WY 2018-2025), and four target ARGs 16 times over two years (WY 2022-2024) when at least one of the two drains was flowing. We hypothesized that nutrient, *E. coli*, and ARG yields exported from the CD field to the recipient ditch would be reduced during managed periods due to reduced drainage flow but would not differ between fields when both drains were unmanaged (i.e., freely draining). We also predicted that DWM would influence *E. coli* and ARG concentrations, with the magnitude and direction of these effects dependent on soil moisture conditions that regulate microbial community dynamics (Austin et al., 2004; Sharma et al., 2019). To our knowledge, this is the first study to simultaneously monitor nutrients, *E. coli*, and ARGs from paired fields under long-term DWM, and one of the few to evaluate DWM performance across multiple years providing an integrated assessment of how drainage management influences biogeochemical and biological contaminant export under variable hydrologic and management conditions.

## 2.0 Methods

### 2.1 Site Description

We conducted this study on working farmland in the Paw Paw River Basin in southern Michigan (United States) during water years (WY; Oct 1-Sept 30) 2017-2025. In 2017, the landowners retrofitted a 13-ha field with pre-existing subsurface tile drainage DWM infrastructure to allow independent control of drainage from two field sections. One section (5.7 ha) was managed with actively controlled drainage during the fallow season (CD), while the other section (7.3 ha) remained free draining (FD) throughout the study period (Fig. S1). On each sampling day, we recorded weir height to document management status. The FD apex height (height from top of DWM box to the top of the weir) was defined at 166 cm. For the CD structure, we set the apex height to 100 cm during controlled periods and 166 cm during free draining periods. These settings mean that we restricted flow during fallow periods and allowed free drainage during the cash crop growing season. Drainage outflows from both CD and FD discharged into an agricultural ditch. Both fields received identical types and application rates of spring-applied synthetic fertilizer through the duration of the study (Table S1). Urea-ammonium nitrate (UAN 28%) was applied three times with an average application rate of 596 L ha^−1^.

Nitrogen was applied three times with an average application rate of 626 L ha^−1^, and calcium and zinc acid amendments were added once (both 9 L ha^−1^). In both spring and fall, equal amounts of liquid dairy manure slurry (Average application 59,937 L ha^−1^) were applied to both sections. In December 2021manure was applied only to the CD section (42,089 L ha^−1^). Although all previous and subsequent manure and fertilizer applications were applied equally to both fields, this management difference may have influenced antecedent nutrient conditions during the following drainage season. The fields were managed under a corn-soybean rotation for the duration of the study.

Soils are predominantly Belleville loamy sand (94.2% of the field area), with smaller extents of Gilford loamy sand (5.3%), and Selfridge loamy sand (0.4%). The 30 yr normal annual precipitation for the site (1995-2025) is 1035 mm yr^−1^. Soil physiochemical characteristics taken in the fall and spring of 2020 and 2021 on each field can be found in Table S2. Fwhile (0-1% slope) and lacks a physical buffer separating the two drainage treatments. Although CD and FD were managed with separate tile drainage systems and independent outlets, subsurface hydrologic connectivity between the fields was not directly evaluated. Therefore, the potential for hydraulic interaction between the systems cannot be fully excluded, although the systems were designed and managed as independent drainage networks. Prior to this study, tile drains were installed at a depth of 76 cm with 30-m spacing throughout most of the field, except in the western portion of the FD section, where spacing was reduced to 15 m to achieve comparable drainage capacity between the FD and CD fields (Fig. S1). The percentage of time the CD field was managed with DWM over each year varied from 50 to 100% (Table S3).

### 2.2 Field Collection

To measure the effects DWM on edge-of-field losses, we took bi-weekly water samples from both CD and FD drains for dissolved nutrients (as NO_3_^−^-N, NH_4_^+^-N, and SRP) from WY 2019-2025, for *E. coli* from WY 2018-2025, and added seasonal water samples for ARGs (*N*=16 dates) from October 2022 to November 2024 (Fig. S2). We selected *E. coli* as a fecal biological indicator because it is a dominant intestinal commensal and potential pathogen commonly associated with livestock manure (Lu et al., 2005). We selected *tetW*, *tetM*, *ermB*, *sul1* as the ARG targets because they represent major antibiotic resistance classes relevant to animal husbandry and human medicine (He et al., 2020). We sampled ARGs 16 times in total, and both drains were flowing on 11 of the 16 sampling dates, allowing paired comparisons of ARG concentrations. ARG yields were compared across all 16 sampling dates. For dissolved nutrients, we filtered 40 mL of tile drain water through type A/E glass fiber filters (Pall, Ann Arbor, MI) into 50 mL centrifuge tubes (Thermo Fisher Scientific, Waltham, MA). We took 300 mL water samples to measure *E. coli* and ARGs using sterile sampling bags (Whirl-Pak, Madison, WI). All water samples were stored on ice until we returned to the lab, after which *E. coli* and ARG samples were processed immediately, and nutrient samples were frozen prior to analysis. On each sampling date, we measured conductivity (μS cm^−1^) using a handheld probe (Cond 3110; WTW), and pH, dissolved oxygen (DO; mg L^−1^), turbidity (Nephelometric Turbidity Units; NTU), and water temperature (°C) using a multi-meter (Multi 3320; Xylem).

We used a continuously deployed pressure transducer (HOBO U20L-0x Water Level Data Logger; Onset Computer Corporation, Bourne, MA, USA) to measure discharge at 15 min intervals. Each time we sampled tile drains, we took hand-measured discharge three times using a graduated bucket and stopwatch. We also deployed a 1 m long SoilVue 10 soil moisture and temperature profile sensor (Campbell Scientific Inc. Logan, UT) in each field (near the tile outlet) with individual sensors placed vertically in the soil profile at five, 10, 20, 40, 50, 60, 75, and 100 cm. The sensors collected data at fifteen-minute intervals from Feb 2021 - Oct 2023. Although very infrequent, the few values (out of over 300,000) that fell outside the possible range of VWC (values less than zero and greater than one) were removed from the dataset from both drains.

### 2.3 Sample Processing and Quantification of E. coli and ARGs

After each sampling event we cultured *E. coli* according to EPA Method 1603 (EPA 2009). Briefly, we vacuum filtered a known volume of water (on average 200 mL) through an MCE membrane filter (Millipore). After filtering we placed on modified mTEC agar (Difco), incubated at 35°C for 2 ± 0.5 hours, then incubated at 44.5°C for 22 ± 2 hours, and counted the target *E. coli* colony forming units (CFU; magenta in color). We analyzed dissolved nutrient samples for NO_3_^−^-N, SRP, and NH_4_^+^-N using cadmium reduction (APHA 2017), ascorbic acid (Murphy & Riley 1962), and phenol-hypochlorite methods (Solórzano 1969), respectively, on a Lachat Flow Injection Auto-Analyzer (Lachat Instruments, Loveland, CO) following standard methods. Instrument detection limits ranged from 2 to 10 µg L^−1^ for NH_4_⁺-N, 2 to 8 µg L^−1^ for SRP, and 0.03 to 0.5 µg L^−1^ for NO_3_ ^−^-N. For values below the detection limit, we substituted one-half the detection limit (US EPA, 2015). To analyze water samples for ARG concentrations, we vacuum filtered each 200 ml sample through a 0.45 μm mixed cellulose ester (MCE) filter (Advantech, Taipei, Taiwan), which we then placed inside a Powerbead extraction tube (Qiagen, Hilden, Germany). We extracted DNA in batches of 24 samples using the DNEasy PowerSoil Pro Kit (Qiagen, Hilden, Germany) following manufacturer instructions and stored both filters and extracted DNA samples at −20 °C until final quantification for ARG targets with digital PCR (dPCR; QIAcuity digital PCR System, Qiagen, Germantown, MD, USA). Our study targets were based on a preliminary screening of the following ARGs: *blaTEM*, *ampC* (enterobacter-specific class C ESBL), *oxa-1*, *int1*, *sul1*, *tetW*, *tetQ*, *mefA*, *ermB*, and *tetM*. We selected *tetW*, *tetM*, *sul1*, and *ermB* because they were all present in the samples, are commonly found in agricultural settings, and are relevant in both human and veterinary health (FDA, 2024) and quantified from total extracted DNA and therefore may include both intracellular and extracellular DNA. We provide complete details for our digital PCR experiments in Supplemental Information (SI; Table S4 and S5).

### 2.4 Continuous load nutrient estimations

We modeled daily nutrient export (as loads; kg day^−1^) from sample concentrations and continuous discharge data using the *loadflex* package in R (Appling et al., 2015). For NO_3_^−^-N and SRP, we selected the composite model, which combines a discharge-concentration regression with interpolation of model residuals to minimize discrepancies between predicted and observed loads. For NH_4_^+^-N, we used linear interpolation because concentration and discharge were not significantly correlated (Vincent et al., 2025). We assessed all models for autocorrelation using the Durbin-Watson test and a first-order autoregressive model (Aulenbach, 2013), and all Loadflex model diagnostics are in the SI. After estimating daily loads, we converted them to yields (kg ha d^−1^) using field area to account for size differences between the two fields. To obtain cumulative discharge (mm), we normalized tile discharge (L s^−1^) by field area and summed over hourly time steps, excluding periods where data were missing from either CD or FD to maintain equal observation counts.

### 2.5 Water yield metrics

We calculated cumulative water yields using only paired observations where both drain sensors recorded valid data. Time steps with missing data due to sensor maintenance or instrument error were excluded from both drain datasets to ensure equivalent comparisons, resulting in the omission of <2% of measurements out of about 136,000 measurements. Data were not normally distributed, so we used a non-parametric Wilcoxon signed-rank test to assess whether percent reductions in annual yields were significant.

To compare mean daily drainage yields between CD and FD, we stratified the dataset into four scenarios (Fig. 1); S1) CD managed with no flow, S2) CD managed, with flow, S3) both drains unmanaged (free flowing), and S4) the entire study period, to assess “net-effect” from 2018-2025. We compared mean daily drainage yield across scenarios S2 to S4 by normalizing discharge by field area, and using a general linear mixed model (GLMM) with drain identity as a fixed effect and year as a random intercept to account for repeated measurements through time using the *glmmTMB* package in R. We specified a Tweedie distribution for unmanaged (S3) and net-effect (S4) scenarios to accommodate right-skewed data containing a mass of zeros, and a Gamma distribution for CD managed with flow periods (S2) when discharge was solely positive. The scenario S1 was not analyzed statistically because discharge from the managed drain was zero during these periods. All model specifications and diagnostics are included in the SI (Table S6). For each drain, we conducted continuous wavelet analyses on daily discharge data using the *waveletComp* package in R to identify dominant flow periodicities and compare variability between CD and FD. Analyses captured periodicities from 1-64 d, spanning seasonal timescales. We normalized wavelet power by series variance and extracted median normalized power across time to summarize variability at each periodicity (Chalise et al., 2023) then compared median power spectra between drains to assess the DWM effect on CD vs FD. To quantify how rapidly the hydrograph rose and fell, we calculated the Richards-Baker Flashiness Index (FI) (Baker et al., 2004) from hourly discharge data by season and water year using the formula:

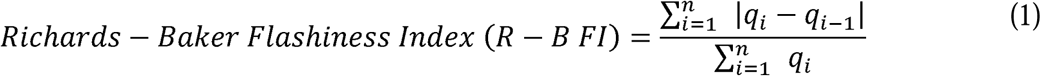

where we used hourly discharge (q) rather than daily discharge to better capture variability in fast-moving subsurface tile drainage systems, departing from the typical calculation approach for flashiness (Deelstra et al., 2014). To compare FI by season, we used a GLMM with a Gamma distribution and log-link, with drain and season as interacting fixed-effects, and WY as a random effect. We used Pearson’s correlation to examine the relationship between annual FI and the percentage of time CD was managed. To complement the analysis of FI, we identified high-flow events as discharge exceeding the seasonal 95^th^ percentile for each drain and compared the number of peaks between drains using a GLMM with drain identity and season as interacting fixed effects and WY as a random effect with a Poisson distribution. All model selection

**Fig. 1.**
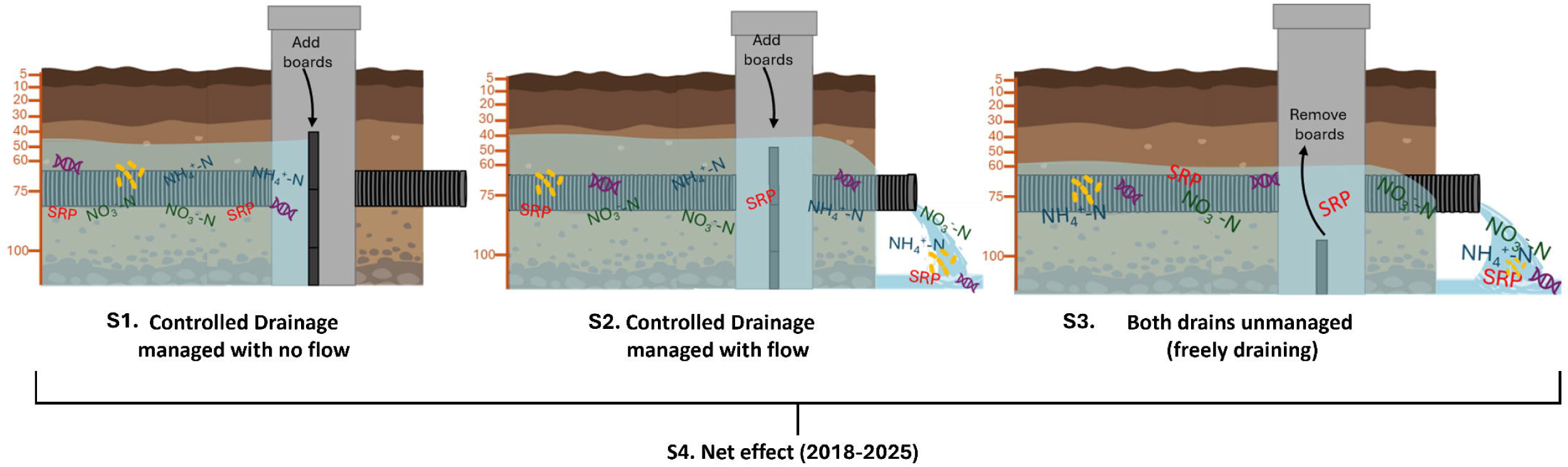
Drainage water management control structures with four different management scenarios: S1) Controlled drainage is managed with no flow, S2) Controlled drainage is managed with flow, S3) Both drains are unmanaged and water can flow freely from the drain, and S4) Net effect, or the entire study period. Measurements from 5-100 cm on the left-hand side of each scenario represent locations of soil moisture sensors (SoilVue 10). Symbols in the water represent antibiotic resistance genes (ARGs; DNA strands), *Escherichia coli* (*E. coli;* microbe cluster), nitrate (NO_3_^−^-N), ammonium (NH_4_^+^-N), and soluble reactive phosphorus (SRP).

### 2.6 Differences in nutrients, E. coli, ARGs, and soil moisture

We compared nutrient, *E. coli*, and ARG concentrations and yields between drains across the three scenarios using the modeling framework described above; all model specifications and diagnostics are described in Table S6. Concentration analyses excluded periods with no flow from CD. Managed-condition comparisons (S2) were restricted to paired observations when both the CD and free drainage (FD) outlets were flowing, whereas the net-effect analysis (S4) combined managed-flow (S2) and unmanaged (S3) periods to represent the overall performance of drainage water management under typical field conditions. For nutrients and *E. coli*, we included drain identity as a fixed effect and year as a random effect, modeling concentrations with a Gamma distribution and log-link and yields with a Tweedie distribution to accommodate zero values (days with no flow from tiles). Analyses were conducted separately for each analyte. For *E.coli* yields during unmanaged periods, adding date as a random effect prohibited model convergence, and was not included (Table S6).

For ARGs, we included date as a random effect and modeled both ARG concentrations and yields using a Tweedie distribution to account for the mixture of zero values and positive continuous concentrations and yields. For all pairwise comparisons, we used estimated marginal means with Bonferroni correction via the *emmeans* package in R. We used the *DHARMa* package in R to evaluate all model fits and inform model selection by assessing residual uniformity, dispersion, and temporal autocorrelation. We report effect sizes as estimated mean ratios (CD relative to FD), obtained from back-transformed emmeans contrasts, and report values in full with 95% confidence intervals in Table S4. These same ratios are expressed in-text as percent change (100 × [estimated mean ratio - 1]).

To compare soil moisture across depths, we subset the dataset to 2021-2023, aggregated 15-minute VWC measurements to monthly means, and used a GLMM to evaluate the effects of drain, depth, and their interaction. Month included as a random effect to account for temporal autocorrelation, with a Gamma distribution and log-link function (Table S6). We also examined whether soil moisture varied with depth by fitting linear regressions between soil moisture and depth by season for each treatment. We used Spearman correlations to assess relationships between ARG concentrations and physicochemical parameters measured on each sampling date, including NO_3_^−^-N, NH_4_^+^-N, SRP, *E. coli*, dissolved oxygen (DO), conductivity, temperature, pH, turbidity, and discharge. We adjusted p-values for multiple comparisons using the Benjamini-Hochberg false discovery rate (FDR) procedure (Benjamini and Hochberg, 1995).

## 3.0 Results

### 3.1 DWM reduced tile flow

Cumulative water yield (mm) from the drain with DWM was significantly reduced (Fig. 2) across all eight study years (Average ± SE; FD: 578 ± 45 mm, CD: 262 ± 44 mm), with a mean reduction of 43% (range: 17-77%; Wilcoxon, p<0.01; Table S7). In addition, mean daily drainage yield (mm) from CD was 56% lower than FD over the study period (GLMM, 95% CI: 52-59%, p<0.0001; Fig. 3), with largest differences during active management periods (S2) when CD had mean flows that were 65% lower than FD (GLMM, 95% CI: 61-69%, p<0.0001; Fig. 3). During unmanaged periods (S3), CD water losses were 14% lower than FD (GLMM, 95% CI: 6-21%, p<0.001).

**Fig. 2.**
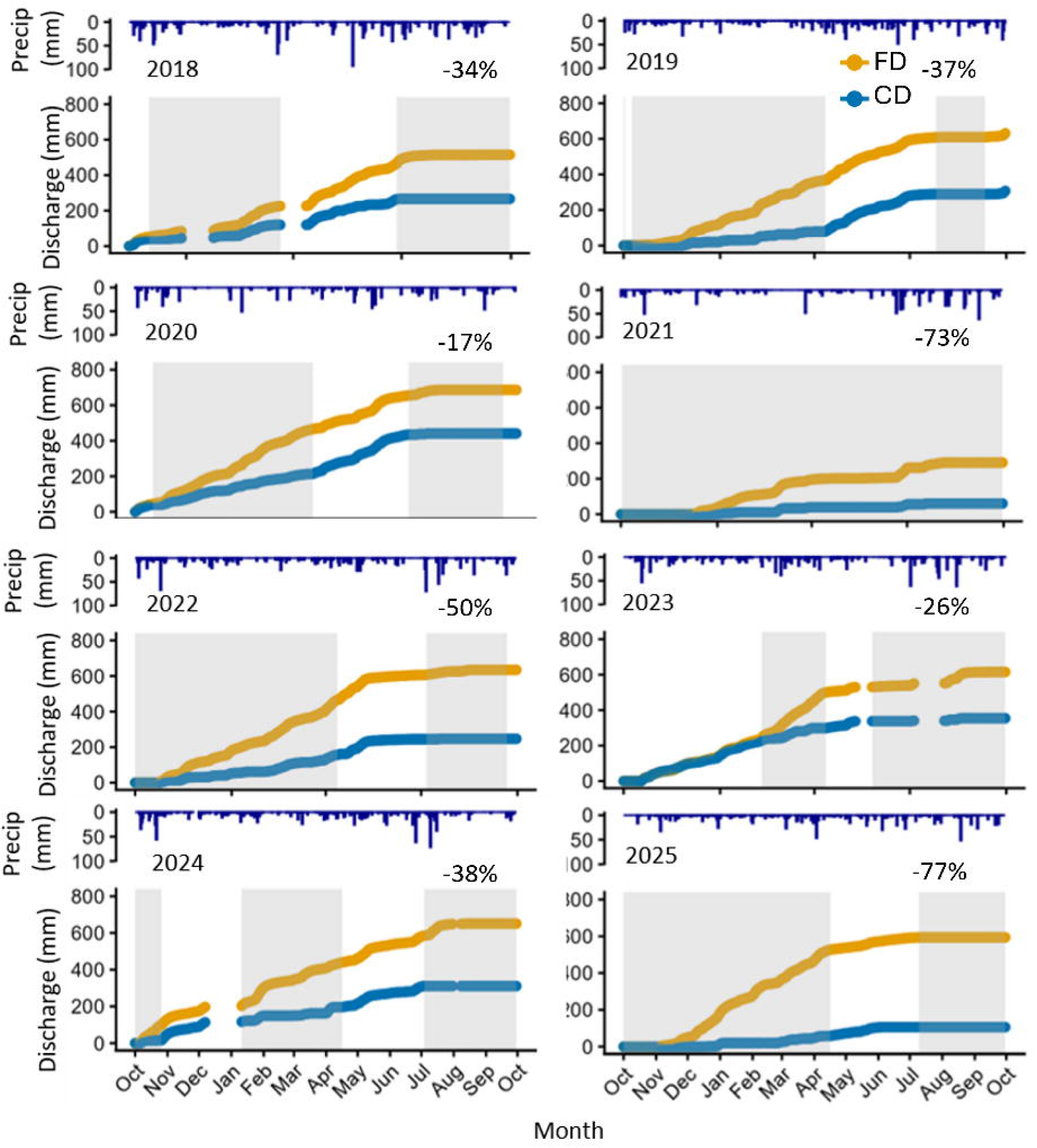
Cumulative discharge (mm) from the controlled drainage (CD; blue) and free drainage (FD; orange) fields over water years (WY) 2018-2025. Precipitation (mm) is indicated as lines on the top of the graphs and shaded rectangles in the background indicate that CD was actively managed during that time. Percent indicates the reduction in cumulative flow in the CD relative to the FD.

**Fig. 3.**
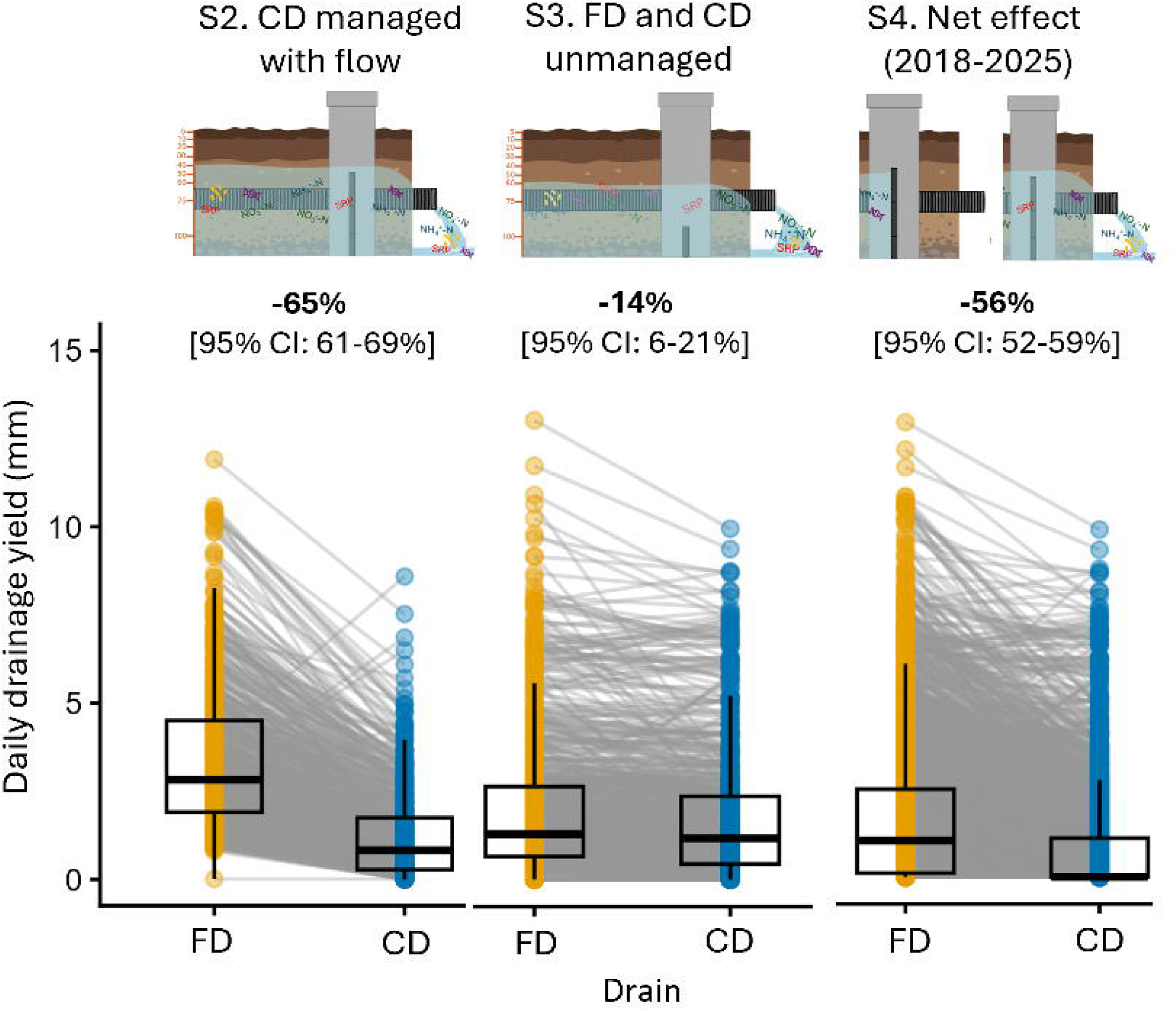
Daily drainage yield (mm) from the controlled drainage (CD) and free drainage (FD) tile drains when CD is managed with flow (S2), both FD and CD are unmanaged (S3), and over the whole study, or “Net Effect” (S4). Percent reduction in daily drainage yield from CD drain is above boxplots with 95% confidence intervals (CI).

Variation in flow, expressed at annual FI, was positively correlated with the percentage of time that CD was actively managed (Pearson’s correlation; r=0.85, p<0.01, Fig. S3a); seasonal FI was higher in CD than FD for every season except fall (EMM, all p<0.05; Fig. S4a). Both drain outlets reached peak flows (top 5%) at similar frequencies across seasons, except during winter when FD flows peaked 2x more often than CD (mean frequency: 8.6 ± 2.2 vs. 3.3 ± 1.0; GLMM, p<0.05). Continuous wavelet transforms revealed strong discharge variability at monthly (∼30 day) and seasonal (∼60 day) timescales in both drains (Fig. S3b), although CD tended to have higher median wavelet power at both monthly and seasonal periodicities (Fig. S3c). Wavelet power was also lower during drier summer months, consistent with reduced flow variability when both drains were more consistently dry.

### 3.2 DWM differentially influenced soil moisture profiles

Sensor data documented changes in soil moisture profiles with and without DWM and soil moisture dynamics differed markedly between treatments, with the direction of the effect reversing with depth (Fig. S5a). From 5-10 cm depth, the CD field had consistently lower mean soil moisture than the FD across all three years (GLMM, all years p<0.05). Deeper into the soil profile, the moisture pattern was reversed and CD had higher soil moisture than FD at 40, 60, and 75 cm. However, soil moisture was similar below the tile line placement at 100 cm (Fig. S5a).

The most pronounced differences in soil moisture between CD and FD occurred at 75 cm, the location nearest the tile line. Controlled drainage at CD maintained higher and sustained soil moisture at 75 cm across all years, and this was consistent independent of individual precipitation events (Fig. S6). In contrast, FD had lower baseline soil moisture punctuated by brief increases following precipitation, after which moisture rapidly returned to near-baseline levels (Fig. S6). These contrasting soil moisture profiles were reflected in distinct depth-moisture relationships for CD compared to FD. Controlled drainage consistently increased mean soil moisture with depth across all seasons (Linear regression, R^2^=0.53-0.85, p<0.001), while there was no significant relationship between depth and soil moisture for FD except for a weak correlation in spring (Linear regression, R^2^=0.09, p<0.05; Fig. S5b).

### 3.3 DWM reduced NO3--N and SRP yields from tiles

Cumulative annual NO_3_ ^−^-N yields from CD ranged from 46-83% lower than for FD (Wilcoxon, p<0.05; Fig. S7a) with a mean reduction of 57.3 kg ha^−1^ per year (Table S7). In contrast, cumulative SRP yields were more variable among years and ranged from 12-82% lower from CD (Wilcoxon, p<0.05) with a mean reduction of 0.20 kg ha^−1^ per year. Finally, cumulative NH_4_^+^-N yields also trended lower from CD compared to FD, but reductions were not statistically significant (Wilcoxon, p>0.05). We also note that NH_4_^+^-N and SRP yields showed step-like patterns in cumulative yields (i.e., more event-driven) compared to the more gradual accumulation pattern of NO_3_^−^-N (Fig. S7b-c). Mean nutrient concentrations did not differ significantly between CD and FD for any solute under any of the drainage management scenarios (GLMM, all p>0.05; Fig. 4). When CD was managed, with flow (S2, Fig. 1), reductions in mean daily yield were significant for all solutes, but largest for NH_4_^+^-N (69%; 95% CI: 38-84%, GLMM, p<0.001), followed by NO_3_^−^-N (62%; 95% CI: 32-78%, p<0.0001) and SRP (52%; 95% CI: 8-75%, p<0.05; Fig. 5). When we considered the entire study period, to assess net-effect from 2018-2025, reductions in mean daily yield were similar and significant for all solutes: NO_3_^−^-N was reduced by 54% (95% CI: 34-67%, p<0.0001), SRP by 50% (95% CI: 23-69%, p<0.05), and NH_4_^+^-N by 48% (95% CI: 12-69%, p<0.05). Yields were not different between the fields during unmanaged periods for any nutrient (GLMM, p>0.05).

**Fig. 4.**
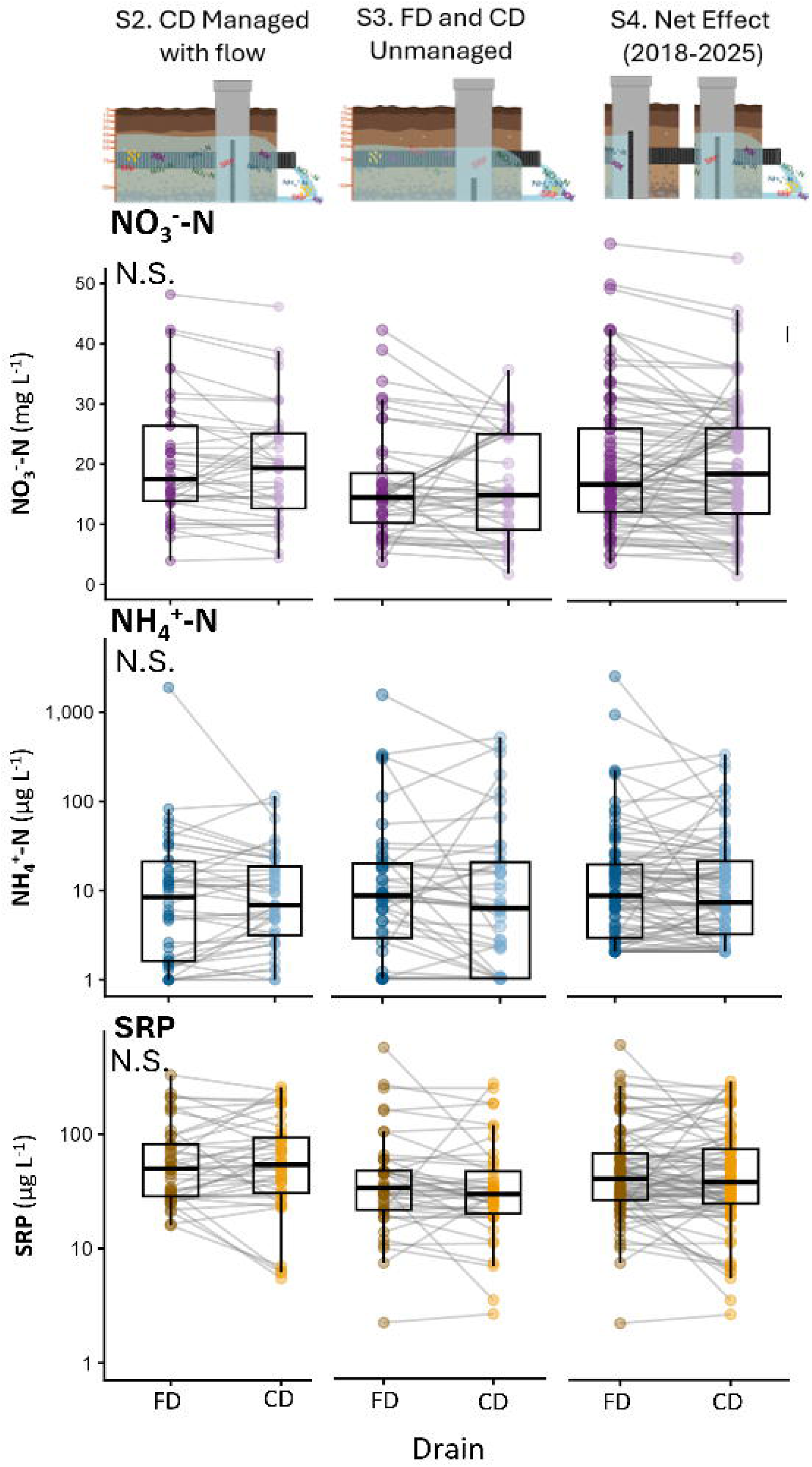
Nutrient concentrations across the free drainage (FD) and controlled drainage (CD) drains across analytes a) nitrate: NO_3_^−^-N (mg L^−1^), b) ammonium: NH_4_^+^-N (µg L^−1^), and c) soluble reactive phosphorus: SRP (µg L^−1^) by three management scenarios: when CD is managed with flow (S2), when both drains are unmanaged (S3), and during the entire study period (Net Effect; S4).

**Fig. 5.**
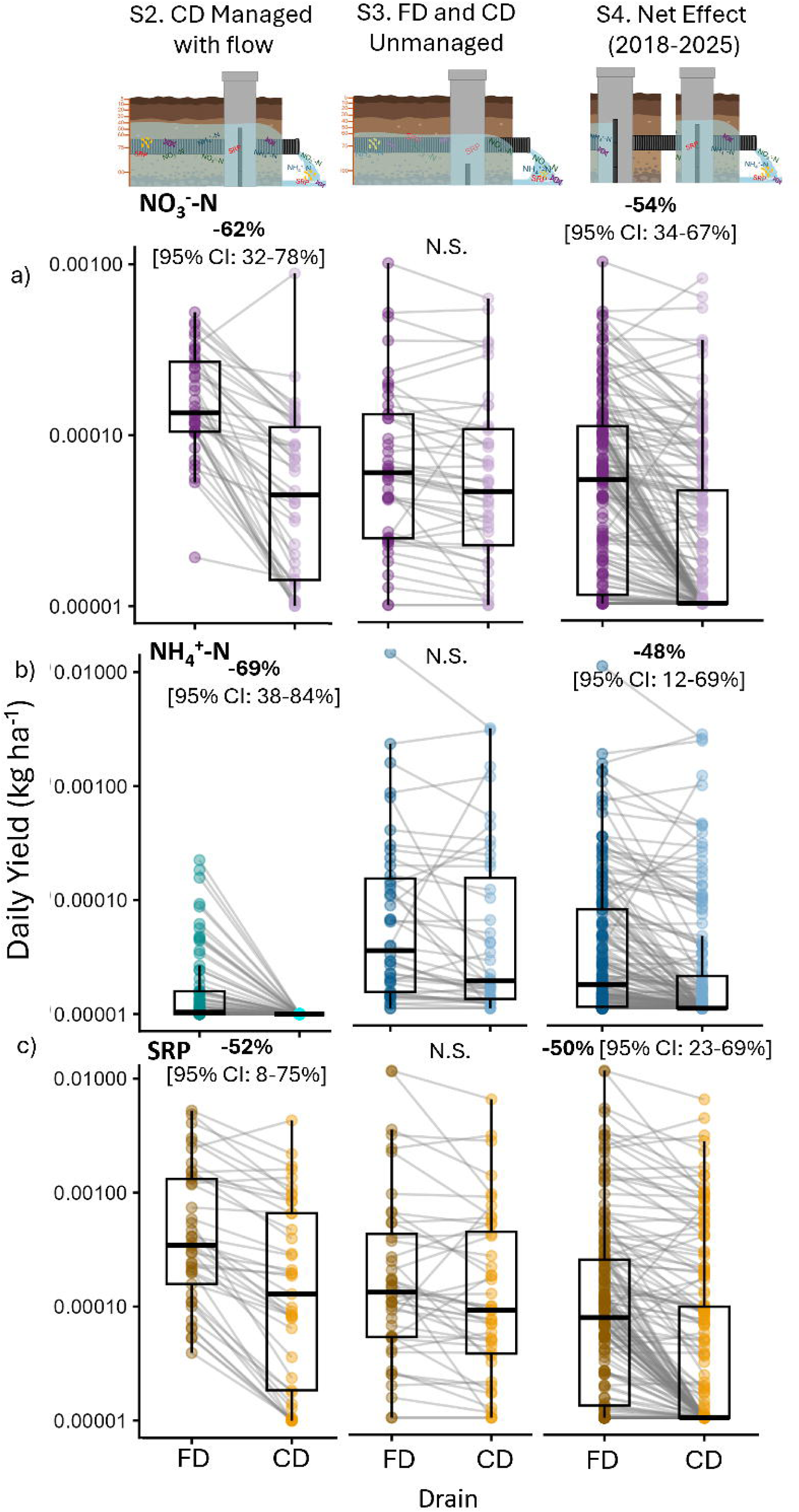
Daily nutrient yields (kg ha^−1^) across the controlled drainage (CD) and free drainage (FD) drains for analytes nitrate; NO_3_^−^-N, ammonium; NH _4_^+^-N, and soluble reactive phosphorus; SRP by three management scenarios: S2) when CD is managed with flow, S3) when both drains unmanaged, and S4) during the entire study period (Net Effect).

### 3.4 DWM reduced E. coli and ARG losses from tiles

Controlled drainage reduced mean *E. coli* concentrations by 46% in CD compared to FD over the full study period (95% CI: 24-62%, GLMM, p<0.001; Fig. 6a), and while concentrations trended lower, concentrations when CD was managed (S2), or both were unmanaged (S3) were not significantly different (GLMM, both p>0.05). Reductions in ARG concentrations were target-dependent: *tetM* and *sul1* were significantly lower from CD under managed (S2; *tetM* = −54%, 95% CI: 43-62%; sul1= −68%, 95% CI: 58-76%, both p<0.0001) and net-effect conditions (S4; *tetM*= −46%, 95% CI: 23-62%; sul1= −58%, 95% CI: 35-73%, both p<0.001), while *tetW* and *ermB* concentrations did not differ significantly between CD and FD under any scenario (GLMM, p>0.05; Fig. 7a). However, a sensitivity analysis excluding a rain-on-snow event on February 28, 2024 (SI Fig. S8, Table S6) indicated that this event strongly influenced observed ARG concentration differences. When this event was removed, the previously observed reductions in *tetM* and *sul1* concentrations under controlled drainage were no longer detected, and no ARG targets showed significantly lower concentrations under controlled drainage across scenarios. Instead, *tetW* concentrations were slightly higher under managed CD conditions (S2; GLMM, p<0.05).

**Fig. 6.**
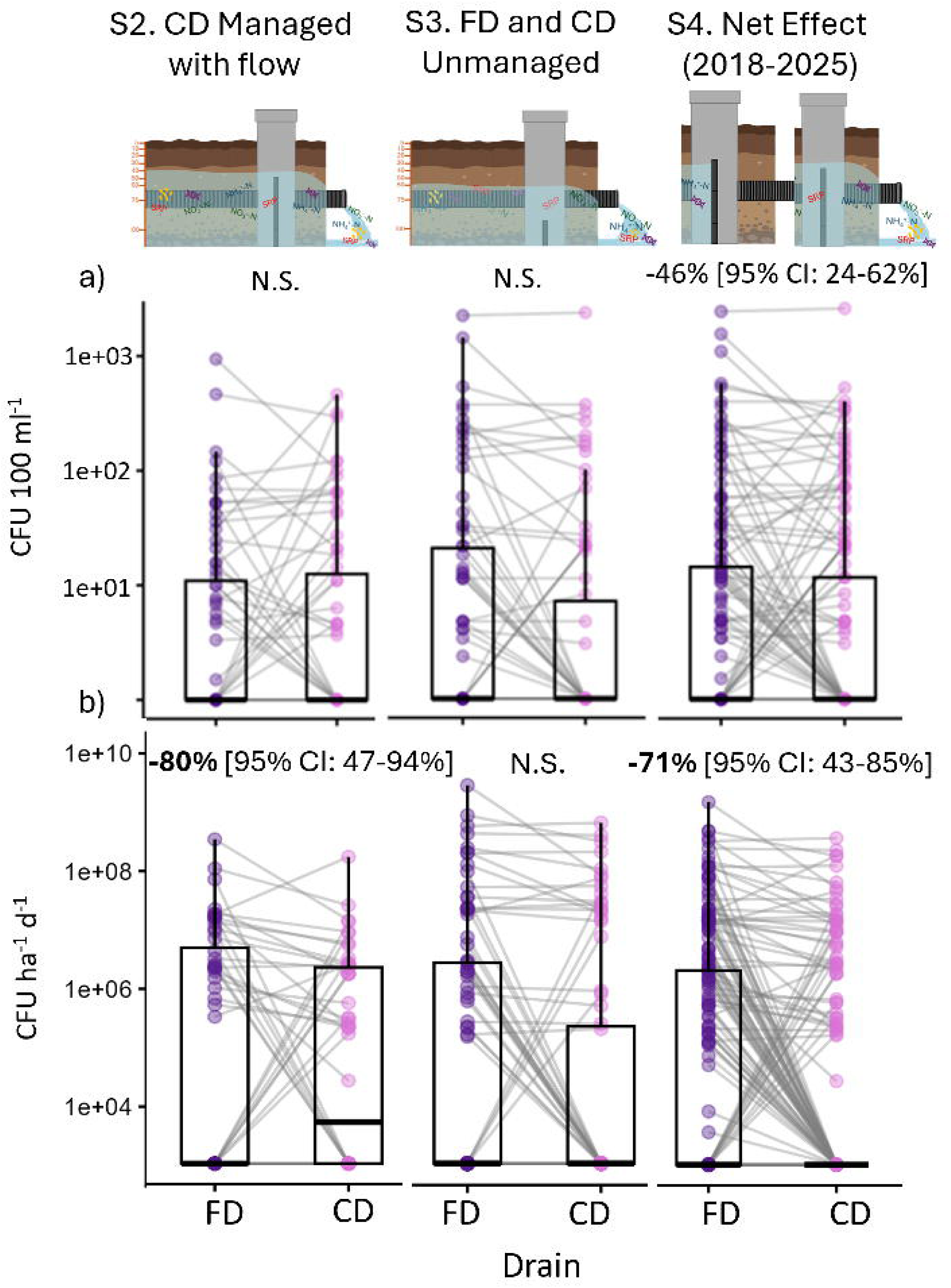
a) *E. coli* concentrations (CFU 100 mL^−1^) and b) *E. coli* yields (CFU ha^−1^ d^−1^) from the free drainage (FD) and controlled drainage (CD) fields across three management scenarios: S2) when CD is managed with flow, S3) when both drains are unmanaged, and S4) during the entire study period (Net Effect).

**Fig. 7.**
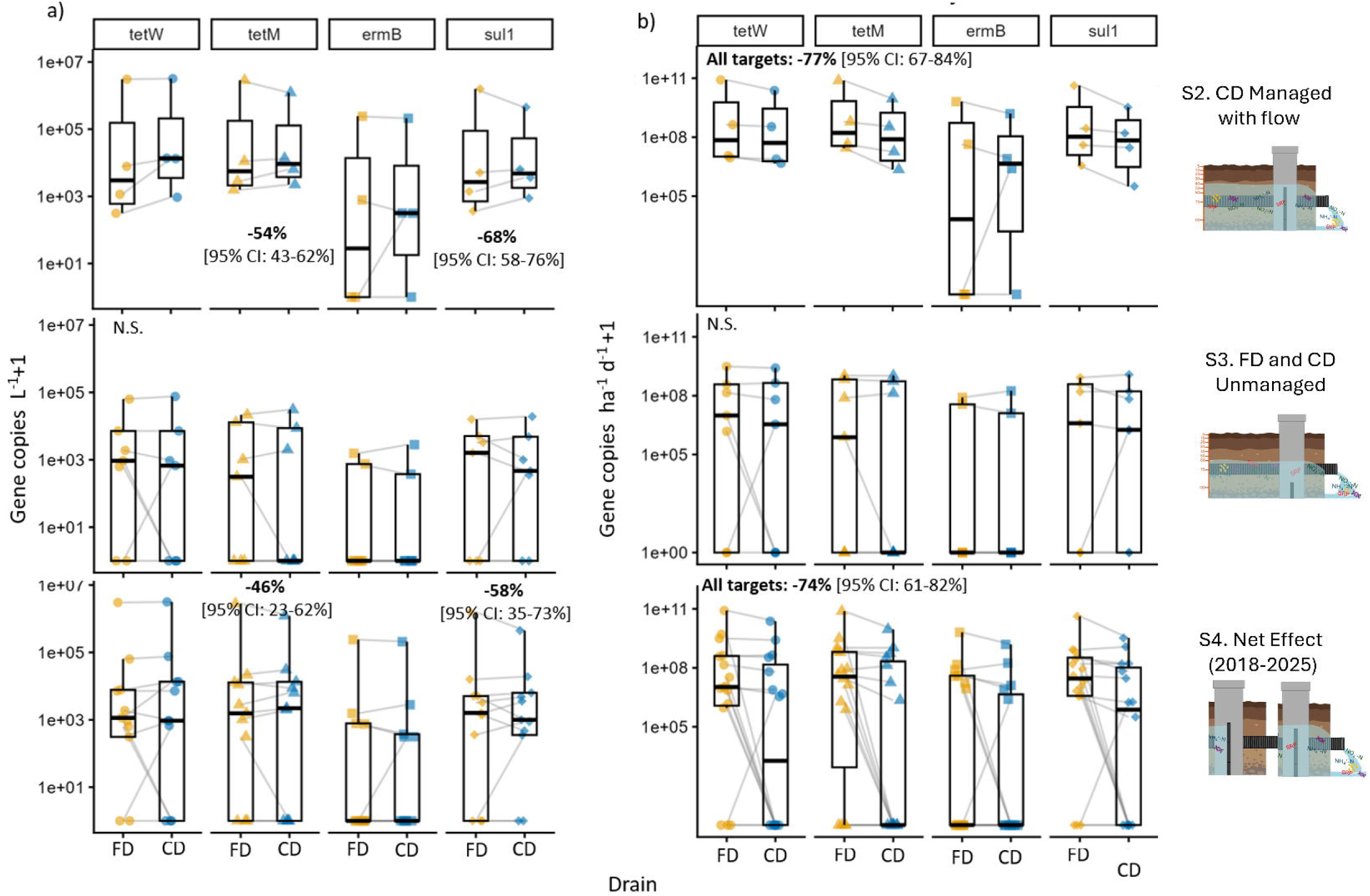
a) ARG concentrations (gene copies liter ^−1^) across the free drainage (FD) and controlled drainage (CD) fields across three management scenarios: S2) CD is managed, S3) both drains are unmanaged, and S4) during the entire study period (Net Effect). Shapes of points indicate different targets, and b) ARG yields (Gene copies ha^−1^ d^−1^+1) across the free drainage (FD) and controlled drainage (CD) drains across three management scenarios: S2) CD is managed, S3) both drains are unmanaged, and S4) during the entire study period (Net Effect).

Controlled drainage reduced mean daily yields for both *E. coli* and ARGs (Fig. 6b; Fig. 7b). When the drain was managed but flowing (S2), *E. coli* yields from CD were 80% lower than FD (95% CI: 47-94%, GLMM, p<0.01), while they were 71% lower across the full study period (S4) (95% CI: 43-85%, p<0.001). Initial analyses indicated that ARG yields were similarly reduced, with CD exporting 77% lower ARG yields during managed and flowing periods (S2; 95% CI: 67-84%) and by 74% over the full study period (S4; 95% CI: 61-82%, both p<0.0001). However, these reductions were strongly influenced by the February 28, 2024 rain-on-snow event. When this event was excluded, the CD drain maintained a 43% reduction in yields during the managed and flowing scenario (S2; GLMM, p<0.0001), whereas the overall net effect (S4) was no longer significant (SI Table 4). These results indicate that while CD reduced ARG export during periods when CD was managed, the magnitude of the full-study reduction was amplified by a single high-flow event. During unmanaged periods (S3), there were no significant differences in ARG concentrations or yields for any target (GLMM, all p>0.05). Lastly, all ARG target concentrations were strongly positively correlated with turbidity (range ρ= 0.61-0.78, p<0.001; Fig. 8). We found that *sul1* concentrations were positively correlated with *E. coli* concentrations (Spearman’s ρ=0.63, p<0.05), and *ermB*, *tetM*, and *tetW* were positively correlated with NH_4_^+^-N concentrations (ρ=0.47, 0.49, 0.57, all p<0.05). Finally, targets *sul1*, *ermB*, and *tetM* were positively correlated with SRP concentrations (ρ=0.60, 0.48, 0.47, all p<0.05). Temperature, DO, conductivity, pH, and discharge did not have a significant relationship with ARG concentrations (p>0.05). Average treatment responses for water yield, nutrients, *E. coli*, and ARGs are included in SI Table S8.

**Fig. 8.**
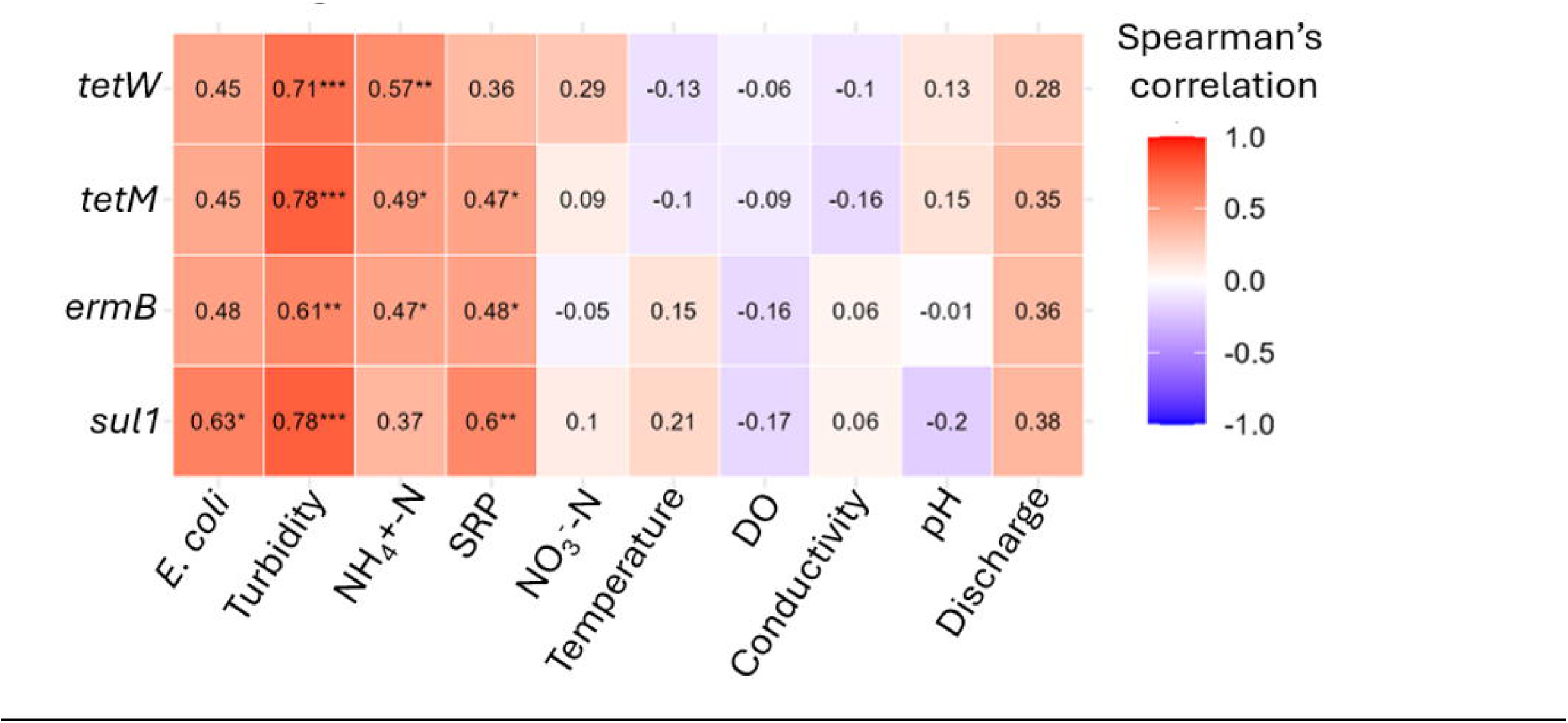
a) Spearman’s correlations between ARG targets *tetW*, *tetM*, *ermB*, *and sul1,* and *E. coli*, turbidity, ammonium (NH_4_^+^-N), soluble reactive phosphorus (SRP), nitrate (NO_3_^−^-N), temperature, dissolved oxygen (DO), conductivity, pH, and discharge (L s^−1^). Stronger positive correlations are red, and more negative correlations are blue. Asterisks indicate statistically significant relationships at α = 0.05, with * = *p* < 0.05, ** = *p* < 0.01, and *** = *p* < 0.001. *P*-values are adjusted for multiple comparisons using Benjamini–Hochberg correction.

## 4.0 Discussion

### 4.1 DWM reduced tile flow

This study provides a novel, multi-year, multi-metric assessment of dissolved nutrients, *E. coli*, and ARG loss reduction under the implementation of DWM. We show that cumulative annual tile flows were reduced from 17-77% when CD was controlled seasonally, which falls within ranges of flow reductions reported in previous studies (Adeuya et al., 2012; Cooke and Verma, 2012; Wesström and Messing, 2007; Williams et al., 2015). In a synthetic review of 31 DWM studies, Frankenberger et al. (2023) found that on average, cumulative flows for CD were 59% lower than for FD tiles (Frankenberger et al., 2023). However, previous work has not frequently isolated the impact of DWM when drainage is managed but still flowing, which occurs when precipitation events increase tile flows and exceed the capacity of the control structure (Skaggs et al., 2012). Our study is among the first to analyze the impacts of DWM on flow when managed but still overtopping the weir, and we demonstrate that DWM can reduce mean daily drainage compared to free flowing tiles, even under high-flow conditions. While we only quantified drainage and solute export from one DWM outlet, future studies should evaluate alternate flow paths and subsurface hydrologic connectivity that may influence observed discharge and contaminant dynamics, as lateral seepage has been identified as a potential contributor to contaminant export from agricultural fields (Lavaire et al., 2017; Liu et al., 2019; Williams et al., 2015).

Additionally, the proportion of time under active CD management varied among years (50.5-99.8%; Table S1), reflecting the operational nature of DWM and variability in hydrologic conditions. Treatment comparisons were based on paired CD and FD observations to prevent periods with longer or shorter management duration from disproportionately influencing treatment effects. However, differences in management duration among years may still influence field conditions (e.g., antecedent soil moisture, drainage connectivity, and water residence times) and affect nutrient and microbial dynamics (Mishra et al., 2008; Outram et al., 2016; Zhang et al., 2026). Therefore, interannual variability in DWM performance likely reflects both differences in management operation and variation in environmental conditions. Hydrologic variability is also an important control on contaminant transport from tile-drained agricultural systems (Osterholz et al., 2024). Metrics such as the Richards-Baker FI quantify rapid fluctuations in drainage discharge that can influence the mobilization and export of solutes (Baker et al., 2004). For example, FI has been shown to explain a large proportion of variability in P leaching across drained agricultural fields, indicating that episodic drainage responses can enhance nutrient transport (Ulén et al., 2016). In our study, annual FI was positively correlated with the proportion of time that CD was actively managed. Despite this increase in FI, cumulative discharge remained substantially lower under controlled drainage, suggesting that DWM reshapes hydrologic patterns while still limiting overall water losses (King et al., 2015; Williams et al., 2015).

### 4.2 DWM differentially modulated soil moisture

moreover We hypothesized that CD would increase soil moisture at all depths relative to the FD field (Yuan et al., 2021); however, the depth-specific results for CD compared to FD were more nuanced. We found that soil moisture at the soil surface (0-5 cm) was lower in CD compared to FD, while soil moisture was higher at depth for CD. We are limited in the comparisons we can make with previous research, because there are few DWM studies published to date that have included soil moisture data. This limits a mechanistic understanding of how tile line depth influences water table dynamics and soil water redistribution, with or without DWM. Moreover, the results are mixed among the studies that have collected soil moisture data using profile-level soil sensors (Duffková et al., 2022; Schott et al., 2017; Woli et al., 2010). Woli et al. (2010) used paired CD and FD fields in Illinois and observed lower VWC in the CD treatment for sensors at 5 and 20 cm, which they attributed to small-scale field effects that outweighed any differences in the water table (Woli et al., 2010). In contrast, Duffková et al. (2022) measured VWC at 20, 40, and 60 cm in paired plots, and found significantly higher soil moisture under DWM at all measured depths (Duffková et al., 2022). Neither study included sensors below 60 cm, leaving the soil moisture regime closer to tile depth (typically at 70-110 cm depth) uncharacterized, which is where we found the largest increases in soil moisture with DWM. Finally, in a range closest to this study, Schott et al. (2017) measured VWC at 10, 20, 40, 60, and 100 cm, and found no significant differences in soil moisture in response to DWM. They attributed the lack of response to a larger timescale that masks shorter-scale fluctuations, and to differences in soil texture (Schott et al., 2017). Although the FD and CD fields were adjacent, agricultural soils are spatially heterogeneous, and localized differences in soil texture and layering have been shown to influence capillary rise and the vertical redistribution of soil water, potentially leading to the shallow layer being drier than anticipated in CD (Li et al., 2014; Woli et al., 2010). Additionally, evapotranspiration has been shown to increase under controlled drainage relative to free drainage (Amatya et al., 1996; Bou Lahdou et al., 2019; Tan et al., 2002), potentially offsetting increases in near-surface soil moisture despite greater water storage deeper in the soil profile. To more fully resolve the effects of DWM on soil moisture dynamics, future work should include thorough pre-DWM soil characterization across multiple paired experimental plots. Additionally, we encourage additional studies that deploy high-frequency soil moisture arrays, covering the full soil depth profile to below the tile line depth. This addition would improve understanding regarding the mechanisms driving nutrient, *E. coli*, and ARG yields in response to DWM.

### 4.3 DWM decreased dissolved nutrient yields

In our study, controlled drainage had the largest impact on average cumulative annual NO_3_^−^-N reduction, followed by SRP, and then NH _4_^+^-N. While NO_3_^−^-N loss reductions due to DWM have been well-documented, fewer studies have evaluated DWM effects on SRP, and this was identified as an area of research need (Ross et al., 2016; Williams et al., 2015). In the Frankenberger et al. (2023) metanalysis, 31 studies measuring NO_3_^−^-N and seven for SRP found that on average, NO_3_^−^-N yields were 46% lower from seasonally managed drains while losses were 34% lower for SRP (Frankenberger et al., 2023). The impact of DWM on NH_4_^+^-N losses is even rarer in paired drain studies, but DWM reduced mean daily NH_4_^+^-N export by 58% in a paired study in Ontario, Canada (Sunohara et al., 2016) and by 28% from a study conducted in rice paddy fields in southern China (Yu et al., 2021). Wang et al. (2020) synthesized 61 controlled drainage studies, ammonium responses were less frequently evaluated than nitrate responses. They found that controlled drainage reduced NH□□-N losses by 18.9% on average but had variable effects on NH□□-N concentrations, highlighting uncertainty in ammonium responses (Wang et al., 2020). Together, our findings demonstrate that DWM reduced nutrient export from tile-drained systems, with reductions largely driven by decreased drainage discharge and the resulting reduction in nutrient mass transport (Skaggs et al., 2012).

Reductions in tile drain losses due to DWM are generally attributed to reductions in flow from tiles, rather than changes in solute concentration (Frankenberger et al., 2023; Williams et al., 2015). In this study, we also partitioned controlled drainage periods into managed periods when the control structure was overtopped and flow continued (S2). Even during managed but flowing periods, mean daily drainage from CD was lower than FD indicating that DWM reduced nutrient yields via its impact on water loss, either by preventing flow entirely (S1) or by slowing drainage during times with more precipitation. This interpretation is supported by the sustained soil moisture that we observed at deeper depths in the CD field, suggesting that controlled drainage delays groundwater recession and reduces drainage following precipitation events compared to the free draining tiles (Dou et al., 2021).

### 4.4 DWM did not change nutrient concentrations

In contrast to yield reductions described previously, no significant differences were found in nutrient concentrations between the CD and FD for any solute. The limited impact of DWM on nutrient concentrations on tile outflows has been well documented previously (Frankenberger et al., 2023; Skaggs et al., 2012; Williams et al., 2015), although a few early studies documented reduced NO_3_^−^-N concentrations with DWM. Kliewer and Gilliam (1995) found that a shallower water table, and increased soil saturation, promoted denitrification in poorly drained soils, thereby reducing NO_3_^−^-N concentrations in tile outflows (Kliewer and Gilliam, 1995). Wesström and Messing (2007) also observed lower NO_3_^−^-N concentrations in spring under DWM, which they attributed to saturated anaerobic conditions (Wesström and Messing, 2007). While we did not assess this, it is possible that soils in our study did not sustain sufficiently anaerobic conditions to stimulate denitrification, as these coarse-textured, sandier soils at this site are less prone to prolonged saturation and oxygen limitation compared to finer-textured soils with higher clay content (Malique et al., 2019).

Previous studies have also raised concerns that DWM may cause soil saturation in the rooting zone and desorption of redox-sensitive P (Grenon et al., 2021); increases in SRP concentrations have been reported (Saadat et al., 2018; Tan and Zhang, 2011), however most research reports no change in SRP concentrations following DWM implementation (Carstensen et al., 2020) which is consistent with our findings. The absence of concentration effects across all three solutes measured here reinforces that mean daily yield reductions observed at this site were driven primarily by reductions in water loss rather than changes in soil biogeochemistry.

### 4.5 DWM reduced both E. coli and ARG yields and concentrations

We found that DWM reduced *E. coli* yields when the CD was managed and flowing, in and over the duration of the study (S2: 80 % and S4: 71% reduction). Field-scale studies of *E. coli* under DWM are limited, but they are generally consistent with yield results reported from this study. Sunohara et al. (2016) reported 76% and 25% reductions in *E. coli* and *Enterococci* spp. yields, respectively, over nine growing seasons (Sunohara et al., 2016). In contrast, watershed-scale responses to DWM appear more variable; Schmidt et al. (2013) observed increased *E. coli* loads in some years under CD, while *Campylobacter* spp. decreased, highlighting the potential variability that is both microbe-specific and driven by changing hydrology among years (Schmidt et al., 2013). The consistency of yield reductions across our eight-year paired field study suggests that field-scale DWM can reduce microbial exports when management is sustained over time.

This study was also the first to document reduced concentrations of *E. coli* over the whole study period. The depth-dependent soil moisture patterns observed under DWM may help explain reductions in *E. coli* concentrations beyond DWM-induced reductions in tile flows. Soil moisture is a key determinant of bacterial survival (Jamieson et al., 2002), and saturated soils have been associated with reduced *E. coli* survival due to limited carbon availability, reduced particle attachment, and increased predation (Chandler and Craven, 1980; Park et al., 2016). Park et al. (2016) reviewed 70 studies and found reduced *E. coli* survival in wet soils compared to intermediate or dry conditions (Park et al., 2016). Underthun et al. (2018) also found peak *E. coli* mortality in sandy loam soils at 25% VWC, with slower decay at 5 - 15% VWC (Underthun et al., 2018). Similarly, Sharma et al. (2019) observed longer *E. coli* survival under lower soil moisture (∼11-12%) compared to higher moisture (∼20%) (Sharma et al., 2019). At our site, the mean VWC at tile depth (∼75 cm) was ∼21% in CD compared to 10% in FD, which means that VWC in CD was approaching the range associated with accelerated *E. coli* mortality in sandy loam soils (Underthun et al., 2018). Elevated saturation near the tile line under DWM may therefore enhance *E. coli* die-off prior to mobilization, contributing to lower concentrations in CD drainage beyond what flow reduction alone would predict.

Initial analysis indicated that DWM reduced ARG yields during managed and flowing periods and during the whole study. Only one previous study has evaluated the impact of DWM on ARG export; which attributed declines to a 67% reduction in flow over 36 days (Frey et al., 2015). For *tetM* and *sul1*, concentrations were also significantly lower during managed and flowing periods (S2; 54% and 68% reductions, respectively), with similar % reductions observed across the full study period (S4; 46% and 58%). That the effect on ARG concentration was not consistent for all targets; *tetW* and *ermB* concentrations did not differ between CD and FD during Scenarios 1, 2, or 4. However, a sensitivity analysis excluding a single rain-on-snow event on 28 February 2024 eliminated this net effect, preserving only differences between the treatments during the managed and flowing periods. These results indicate that ARG export was strongly influenced by this hydrologic event. Studies have shown that ARG export into nearby watersheds is highly affected by even moderate intensity precipitation events (Di Cesare et al., 2017; Jamieson et al., 2002). Sampling in this experiment was designed to capture at least two events per season to characterize differences in ARG abundance associated with agricultural management. Given this temporal resolution, ARG results may be more susceptible to seasonal variability and event-specific conditions (e.g., precipitation timing, manure or fertilizer application, management changes) occurring between sampling dates. These findings suggest that extreme hydrologic events may disproportionately influence ARG mobilization and transport, emphasizing the importance of multi-year sampling that captures a range of hydrologic conditions.

Variation in concentrations among the four ARG targets may reflect differences in host range and form in which they persist in the environment (Zhang et al., 2009). We note that ARG genes such as *sul1*, *ermB*, *tetM*, and *tetW* occur across diverse bacterial hosts in mixed microbiomes (Agga et al., 2020; Tsiklauri et al., 2025; Wang and Chai, 2022). Elevated soil moisture under DWM may promote anaerobic conditions that support persistence of bacterial hosts adapted to these conditions (Jamieson et al., 2002). Anerobic conditions may reduce interactions between ARGs and mobile genetic elements that contribute to horizontal gene transfer, promoting the dissipation from the environment (Xu et al., 2021). However, moist environments can also foster microbial reproduction and ARG dissemination (Brockett et al., 2012; Zhu et al., 2022). Additionally, ARGs can exist as extracellular DNA (eDNA) and biotic and abiotic environmental characteristics can influence their prevalence in various environments (Zarei-Baygi and Smith, 2021). For example, eDNA has a high binding ability and can be adsorbed by soil and particles (Poly et al., 2000; Mao et al., 2014). In this study, all ARG targets were highly correlated with turbidity, however, average differences in turbidity between drainage treatments were small (Average NTU; CD= 5.4 6.6, FD=5.8 7.3), suggesting that reduced ARG export was not solely attributable to reduced sediment transport. Although the effects of soil moisture on ARG adsorption kinetics remain largely unknown, research on other contaminants suggests that changes in soil moisture can influence sorption processes and the retention of compounds and particles within soil matrices (Ashraf et al., 2024; Zarzsevszkij et al., 2024). Future studies should investigate ARG bacterial host associations in systems with DWM, and link redox conditions and soil moisture conditions with gene persistence and transport.

An additional limitation is that this study was conducted using a single paired field design. Although the controlled drainage CD and free drainage FD fields were managed under the same cropping system and monitored over multiple years, field-scale characteristics, like soil heterogeneity, drainage configuration, and local management history likely influence the effectiveness of DWM (Ross et al., 2016; Skaggs et al., 2012). Consequently, the magnitude of nutrient, *E. coli*, and ARG reductions reported here may not be directly transferable to other tile-drained agricultural systems. The measurement of these three contaminant classes across additional sites encompassing a broader range of soils, climates, and management practices on working farmlands will be important for evaluating the generality of these findings in realistic use-cases.

## 5.0 Conclusion

Overall, microbial and nutrient reductions in runoff as a result of DWM were decoupled, suggesting that controlled drainage influenced contaminant export through mechanisms that extend beyond flow reduction alone, and confirming that reductions in dissolved nutrient yields are driven primarily by decreases in tile flow. Novel recorded reductions in *E. coli* and concentrations for two of four ARGs suggest additional biological mechanisms may also be occurring, however ARG dynamics were heavily influenced by a large precipitation event. Our results suggest that the impact of DWM on nutrient and biological contaminants are complex and context-dependent, likely varying with soil type, management intensity, variations in the water year, and specific contaminant characteristics. Future work examining microbial community composition, ARG-host relationships, and soil redox dynamics across depths and seasons would help clarify the mechanisms driving these patterns. While DWM was not designed to reduce microbial determinants, the event-driven reductions in ARG concentrations, and consistent *E. coli* concentration reductions across eight years suggest that biological contaminants warrant consideration alongside nutrients in future management decisions. This study represents one of the first long-term assessments of the impact of DWM on NO□□-N, NH□□-N, and SRP export from working agricultural fields, and the first to evaluate these nutrients alongside E. coli and ARGs. High-frequency soil moisture sensors provided mechanistic insight into treatment effects, while long-term, high-frequency monitoring strengthened our ability to quantify changes in nutrient and microbial exports across a range of hydrologic conditions. Our findings demonstrate that existing nutrient management practices may provide co-benefits beyond nutrient reduction by mitigating *E. coli* and ARG export. These results highlight the potential for controlled drainage and other drainage water management practices to be considered within broader agricultural conservation programs, watershed planning efforts, and future management practices that address microbial contamination and antimicrobial resistance alongside traditional nutrient pollutants. Incorporating microbial outcomes into evaluations of water quality management strategies may improve the effectiveness of interventions designed to protect downstream water resources.

## Acknowledgements

This research was supported by grants from the EPA Great Lakes Restoration Initiative (#GL-00E02207), the Great Lakes Protection Fund (Project #1212), Environment, Great Lakes, and Energy grants #2024-0020 and #2020-0028, and the USDA Grant #1024735. We would like to thank our partners at the Van Buren County Soil and Water Conservation District, and to our collaborators at Xylem. We would also like to thank the farmers and landowners for providing access to their land for the duration of the study. Thank you to the many Tank lab members that assisted with data collection and analysis including Dr. Anna Vincent, Dr. Abagael Pruitt, and Dr. Elise Snyder. Thank you to the many Bibby lab members for their assistance in the field and laboratory including Dr. Olivia Ginn, Dr. Bethany Oceguera-Johnson, Xijiao Wang, Sushil Dahal, Misa Muranaka, Patrick Kneubler, and Caleigh Dahn.

## CrediT authorship contribution statement

**Emma M. Thrift-Cahall:** Writing- original draft, Visualization, Methodology, Investigation, Formal analysis, Conceptualization, Project administration, Data curation. **Jennifer L. Tank:** Methodology, Writing- review & editing, Supervision, Project administration, Investigation, Funding acquisition, Formal analysis, Conceptualization, Resources. **Ursula H. Mahl:** Writing-review & editing, Data curation, Project administration, Supervision, Formal analysis, Methodology**. Mitchell J. Liddick:** Investigation, Writing- review & editing. **Andrei Badilla-Aguilar:** Data curation, Investigation, Writing-review & editing. **Kyle Bibby:** Writing- review & editing, Supervision, Project administration, Investigation, Funding acquisition, Methodology, Conceptualization, Resources.

## Data availability

The data generated and analyzed in this study will be made publicly available on GitHub (https://github.com/ethriftcahall) upon publication.

