## Supplementary Material for "Drainage water management reduces nutrient, *Escherichia coli*, and antibiotic resistance gene export"

**Supplemental Information**

**Text**

1. Loadflex model diagnostics
2. dPCR specifications

**Figures**

1. S1-S8

**Tables**

1. S1-S8

**Loadflex model diagnostics:**

For model selection, we selected solute-specific models based on the strength of discharge-concentration relationships and residual diagnostics. For free drainage (FD), we selected the NO_3_⁻-N composite model because it reduced residual autocorrelation compared with regression-only approaches (Durbin-Watson = 1.97; ρ = 0.02). We selected the NH_4_⁺-N interpolation model because discharge explained little variation in concentrations (Linear Model: R^2^ = 0.08), and regression-based models did not improve model performance. The SRP composite model was selected because SRP concentrations exhibited a significant relationship with discharge (R^2^ = 0.16, p < 0.0001), and the composite model minimized residual autocorrelation (Durbin-Watson = 2.03; ρ = -0.02). For CD, we selected the NO_3_⁻-N composite model because it reduced residual autocorrelation compared with regression-only approaches (Durbin-Watson = 1.79; ρ = 0.10). We selected the NH_4_⁺-N interpolation model because discharge explained little variability in concentrations (Linear Model: R² = 0.07, p < 0.05), and regression-based approaches provided limited improvement over direct interpolation of observed concentrations. We selected the SRP composite model because SRP concentrations exhibited a significant relationship with discharge (Linear Model: R^2^ = 0.13, p < 0.01), and the composite model minimized residual autocorrelation (Durbin-Watson = 2.31; ρ = -0.16).

**dPCR specifications:**

We assayed all DNA extracts using previously developed primer and probe combinations for *tetW* (Knapp et al., 2009), *tetM* (Peak et al., 2007), *ermB* (Böckelmann et al., 2009), *sul1* (McKinney et al., 2018). The reaction mix contained 2 µL of template, a primer concentration of 800 nM, a probe concentration of 400 nM, and 1X QIAcuity Probe Mastermix (Qiagen), for a total volume of 14 uL. For all assays on each dPCR plate, we included a positive template control and negative template control. We performed manual thresholding for each target in the QIAcuity Software Suite and maintained the target-specific threshold levels for the whole experiment. We determined the 95% limits of detection (LODs) for *tetW* and *ermB* experimentally, using 10 replicate serial dilutions of positive controls and a probit analysis as outlined by Stokdyk et al. (2016) (Stokdyk et al., 2016). For *tetM*, samples containing >2 positive partitions were considered 'positive' (Qiagen guidelines: <https://www.qiagen.com/us/applications/digital-pcr/beginners/dpcr-guide/setup-and-troubleshooting>)

**Figures**

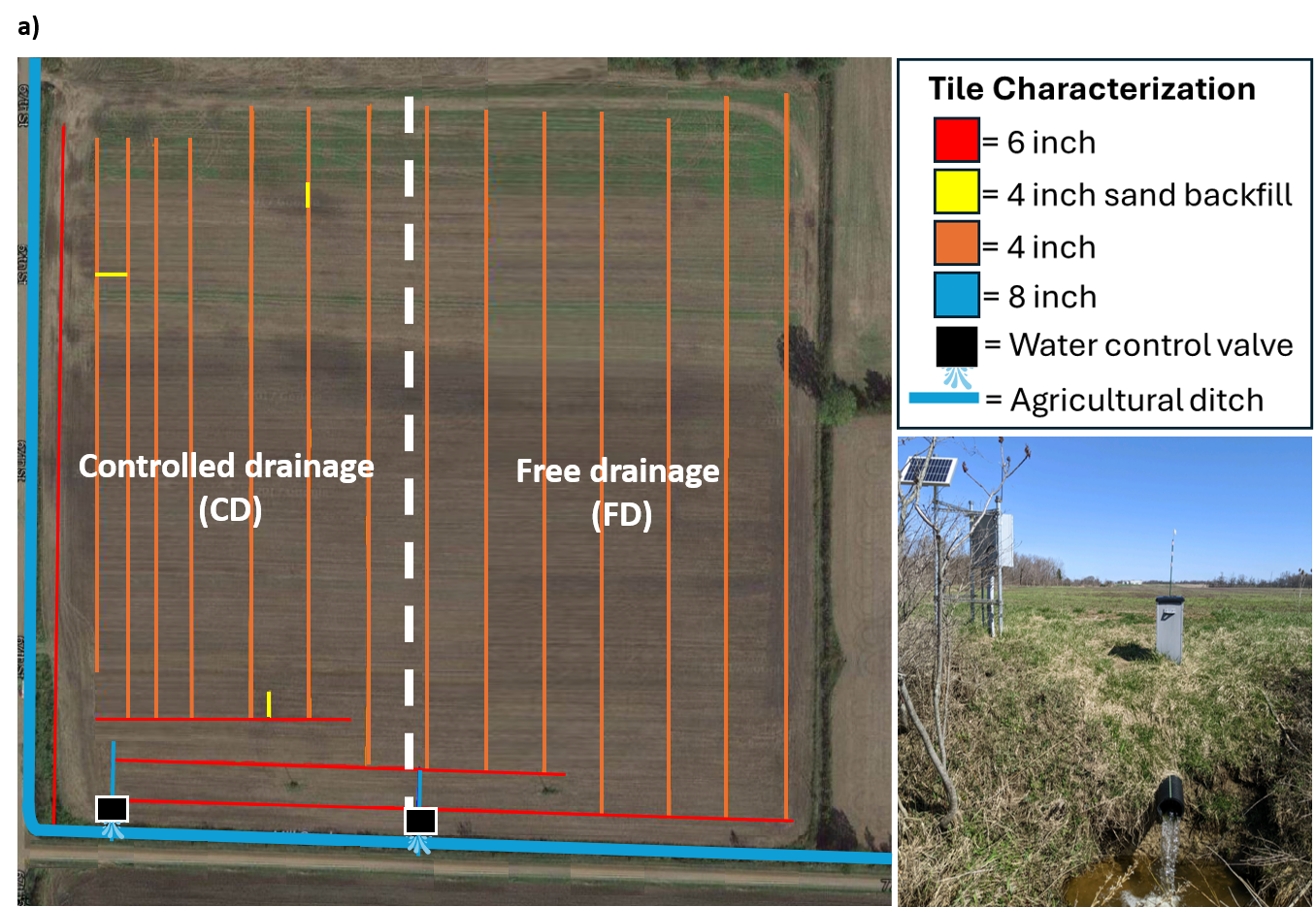

**Fig. S1** a) Tile drainage lines in fields with paired management. The controlled drainage (CD) field has a Drainage Water Management (DWM) structure installed to manually adjust flow, and free drainage (FD) field remains free flowing. Tile drains with different diameters are indicated by different colors. b) DWM structure draining into the agricultural stream.

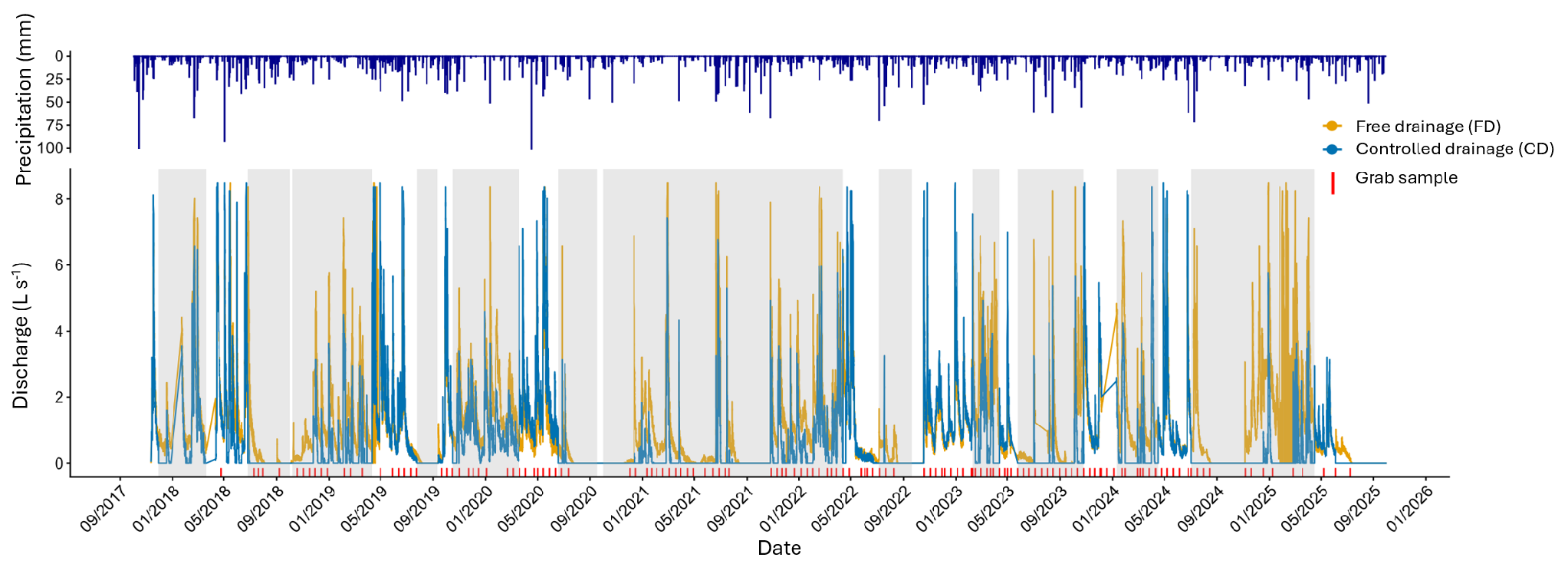

**Fig. S2** Hydrograph of discharge (L s^-1^) from the controlled drainage (CD; blue), and the free drainage (FD; orange), with precipitation (mm) shown above. Dates of grab samples are indicated in red; sampling days when tile drains were not flowing are not shown. Shaded regions indicate time periods when CD was actively managed.

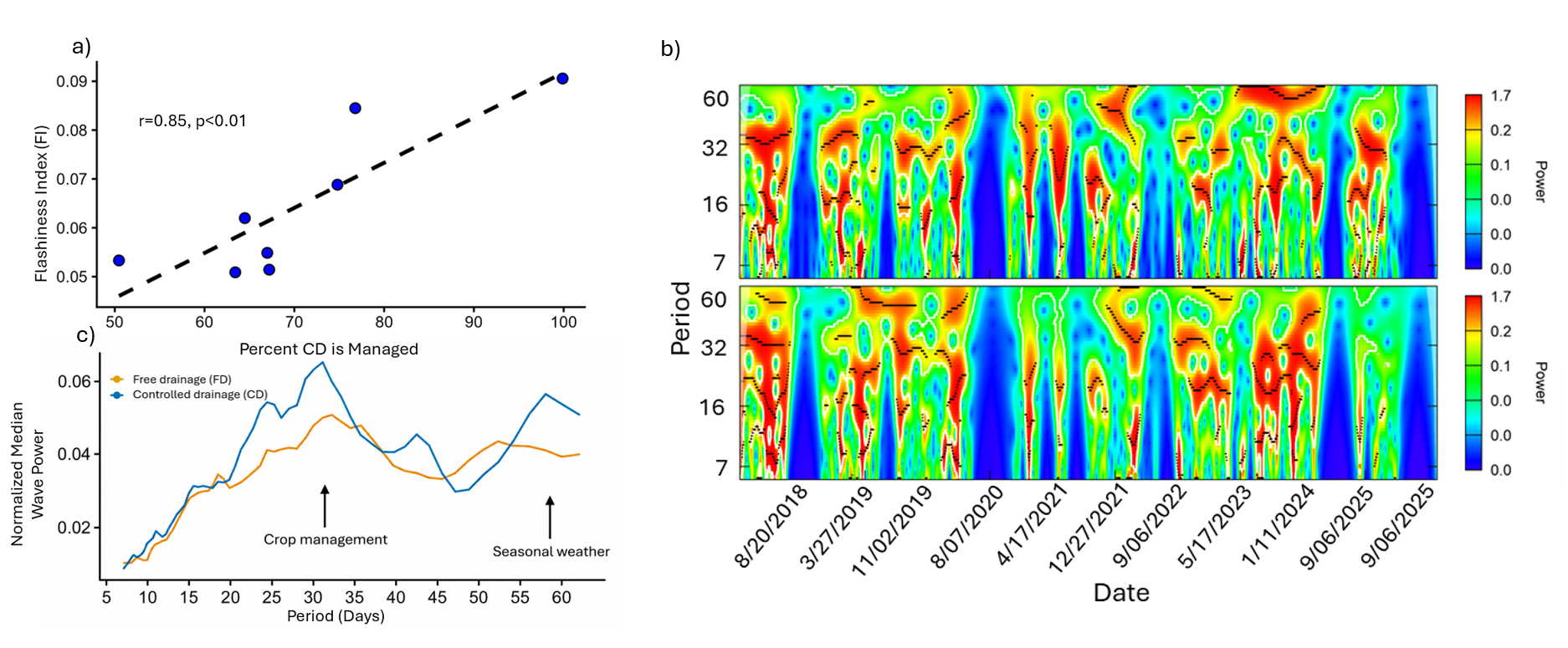

**Fig. S3** a) Linear regression between Flashiness Index (FI) and the percent the controlled drainage (CD) outlet is managed, b) wavelet transform spectra, with color indicating wavelet power, period on the y-axis, and date on the x-axis, c) normalized median wavelet power for free drainage (FD; orange) and controlled drainage (CD; blue) across periods. Arrows indicate examples of crop management activities and seasonal weather patterns that may occur at these timescales.

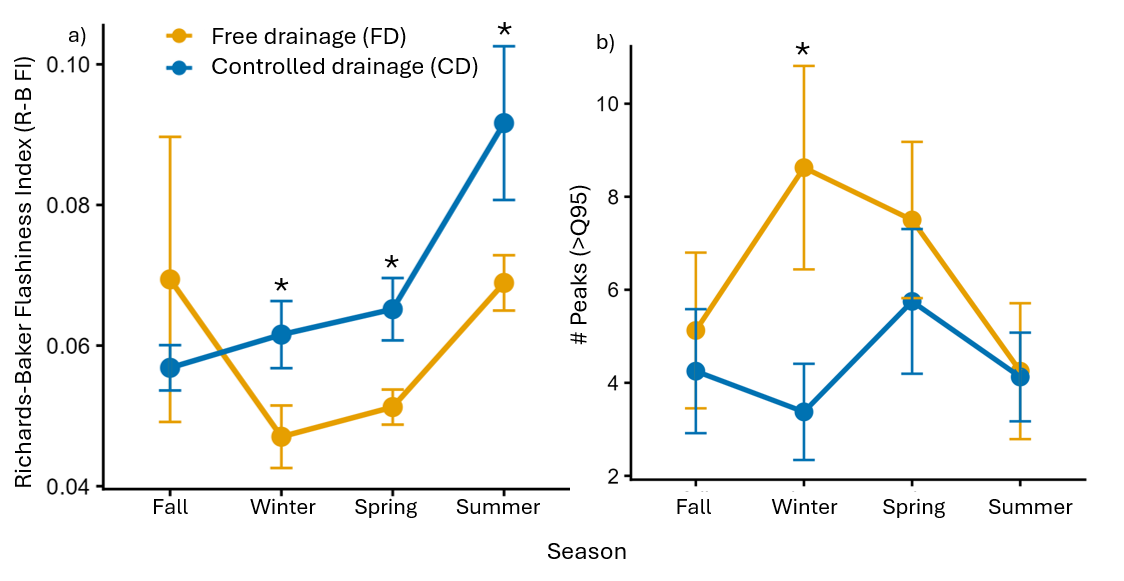

**Fig. S4** a) Average Flashiness Index (FI) across seasons; b) number of discharge peaks that exceed 95% of total flows by season. Orange dots represent mean values for the free drainage (FD) field, and blue dots represent mean values for the controlled drainage (CD) field ± standard error (SE). Asterisks indicate statistically significant differences between paired drains.

**Fig. S5** a) Average monthly soil moisture (volumetric water content; VWC), across soil profile depths (5, 10, 20, 30, 40, 50, 60, 75, and 100 cm) with lines connecting paired values across the same months and years (2021-2023). Percent reduction or increase in soil moisture across fields is indicated above the boxplots with corresponding confidence intervals (CI); N.S. indicates no significant difference. b) Linear regression between VWC and soil depth across seasons by drain. Regression lines indicate that the relationship is significant (p<0.05) and R^2^ represents the
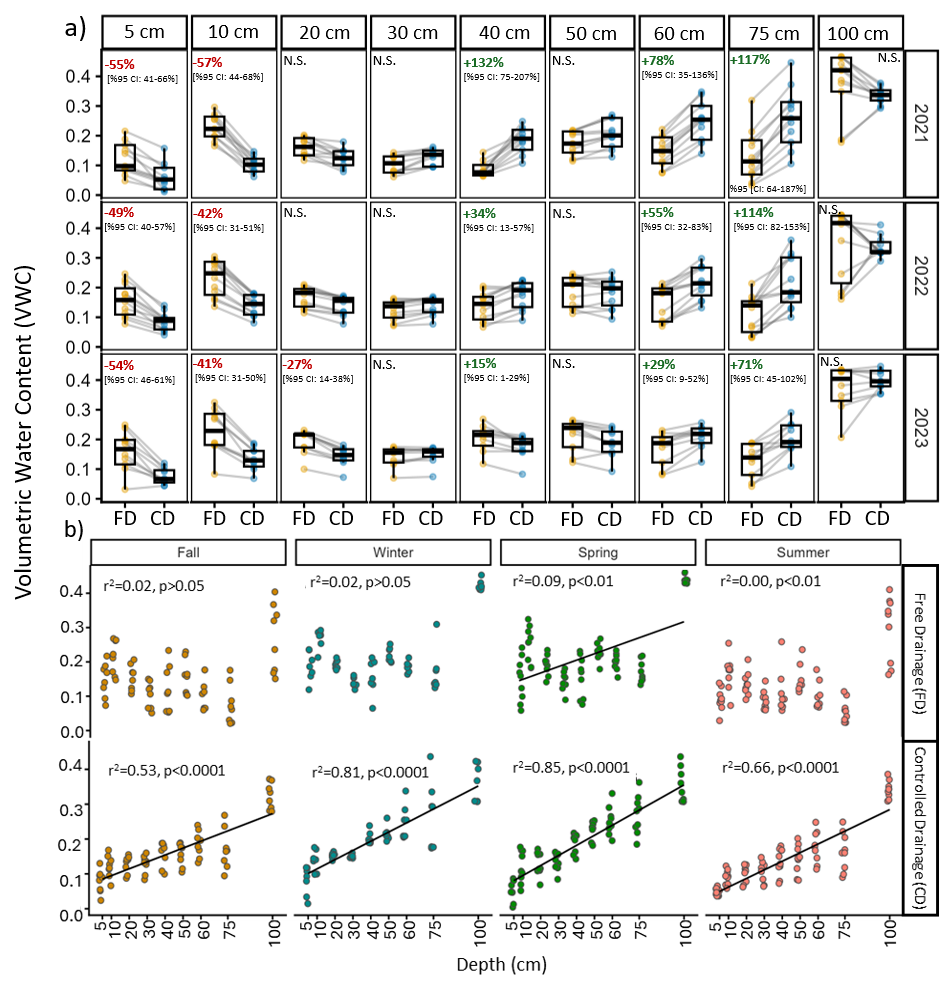
coefficient of determination.

**Fig. S6** Soil moisture (volumetric water content; VWC) at 10, 20, 40, 60, and 75 cm below the soil surface over the period of record (02/2021-12/2023). Shaded background indicated periods in which the controlled drainage (CD) field was managed. Orange lines represent free drainage (FD) soil moisture, and teal lines represent CD soil moisture. We removed periods with missing soil moisture data or data outside the limit of possibility for both drains if data were missing in one drain to ensure equitable comparison. For visualization purposes, we smoothed volumetric water content using a 7-day centered rolling mean within each field and depth
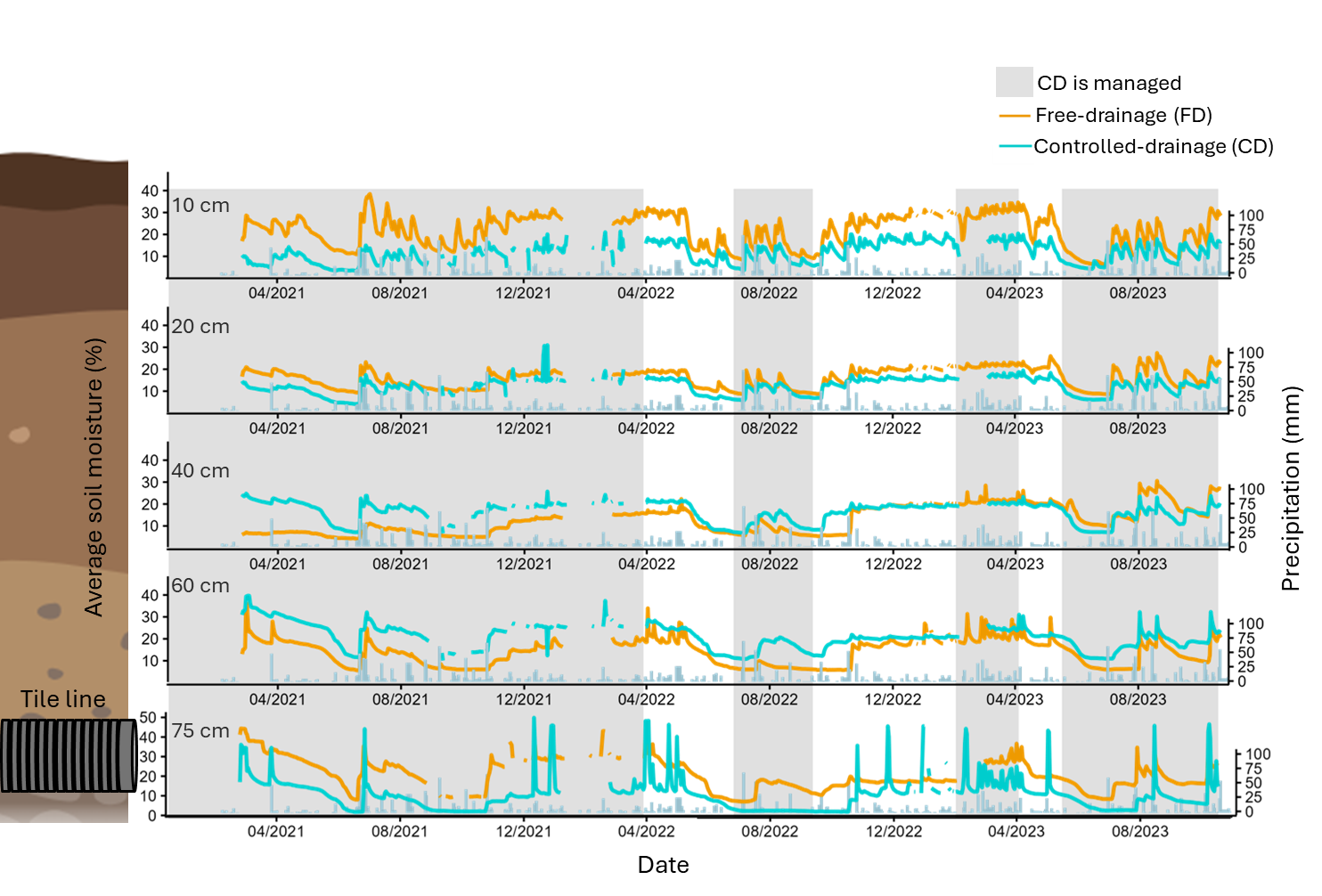
.

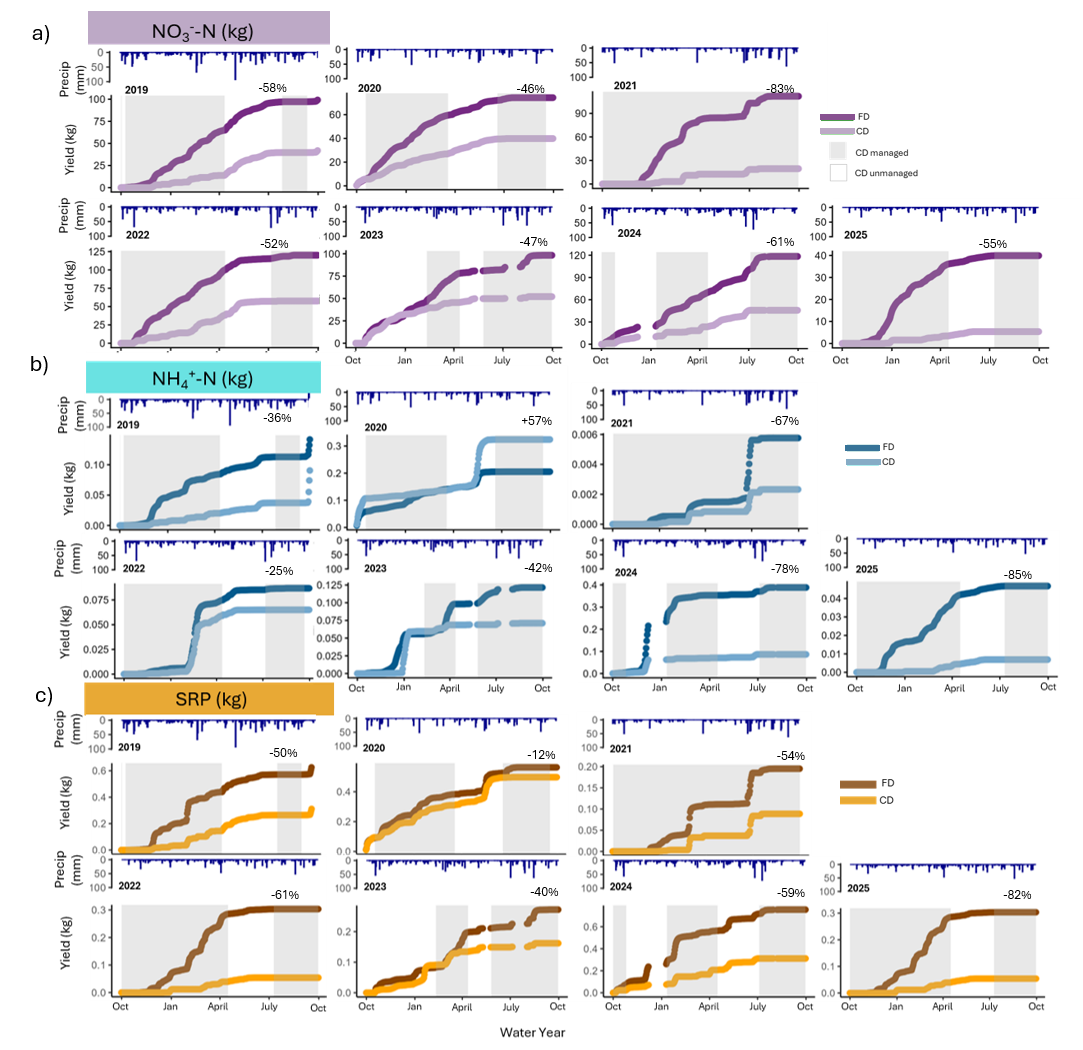
**Fig. S7** Cumulative nutrient yields (kg) by water year from Loadflex for a) nitrate (NO_3_^-^-N), b) ammonium (NH_4_^+^-N), and c) soluble reactive phosphorus (SRP). The darker colors represent the free drainage (FD), while lighter represent controlled drainage (CD). Precipitation (mm) is layered on top, with shaded background rectangles indicating periods when CD was actively managed. Percent reductions are included above the lines, and reference drainage from CD compared to FD.

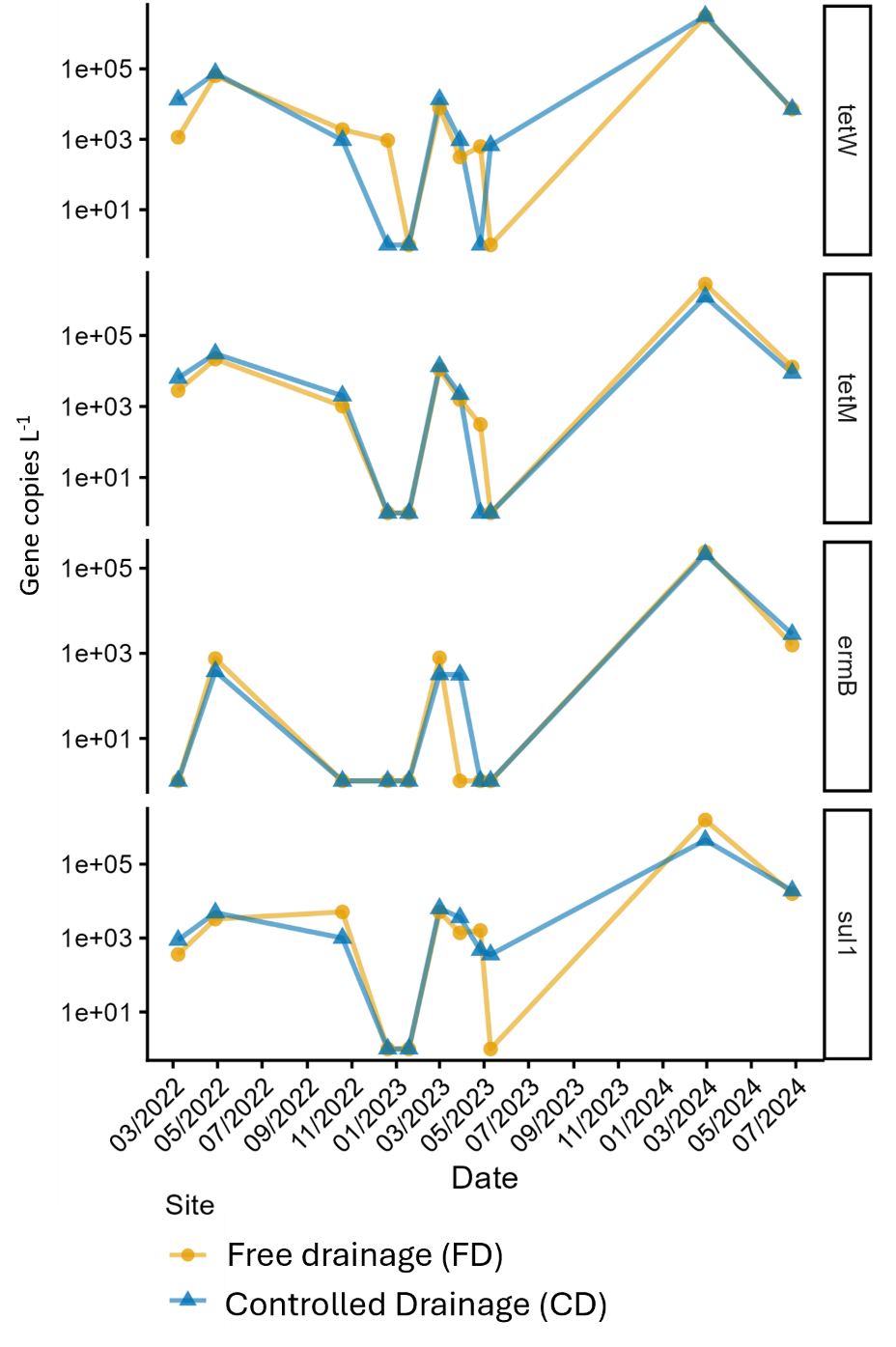

**Fig S8** Time series of antimicrobial resistance genes (ARG) concentrations (gene copies L^-1^) from across sampling dates. Orange circles represent values from free drainage (FD), and blue triangles represent values from controlled drainage (CD).

**Tables**

**Table S1** Field application rates and timing of fertilizer, manure, and other amendments.

| **Date** | **Field zone** | **Field practice** | **Type** | **Application rate (if applicable)** | **Units** |
| --- | --- | --- | --- | --- | --- |
| 6/7/2018 | Both | Fertilizer | UAN 28% | 398.44 | L ha^-1^ |
| 10/3/2018 | Both | Manure |  | 28059 | L ha^-1^ |
| 9/19/2019 | Both | Manure |  | 28059 | L ha^-1^ |
| 6/3/2020 | Both | Fertilizer | UAN 28% | 392.83 | L ha^-1^ |
| 7/3/2020 | Both | Fertilizer | UAN 28% | 996 | L ha^-1^ |
| 9/24/2020 | Both | Manure |  | 16873 | L ha^-1^ |
| 12/9/2021 | CD | Manure |  | 42089 | L ha^-1^ |
| 5/12/2022 | CD | Manure |  | 87198 | L ha^-1^ |
| 5/18/2022 | FD | Manure |  | 87198 | L ha^-1^ |
| 6/1/2022 | Both | Fertilizer | Nitrogen | 41 | kg ha^-1^ |
| 7/1/2022 | Both | Fertilizer | Nitrogen | 97 | kg ha^-1^ |
| 7/1/2022 | Both | Fertilizer | Calcium | 9 | L ha^-1^ |
| 11/16/2023 | Both | Manure |  | 112236 | L ha^-1^ |
| 5/31/2024 | Both | Fertilizer | Biosystem product Zinc acid (Zinc, sulfur, and organic acids) (4-6-4) | 9.35 | L ha^-1^ |
| 7/3/2024 | Both | Amendment | Biosystem product Zinc acid (Zinc, sulfur, and organic acids) (4-6-4) | 9.35 | L ha^-1^ |
| 7/3/2024 | Both | Fertilizer | Nitrogen | 98 | kg ha^-1^ |
| 5/12/2025 | Both | Manure |  | 78107 | L ha^-1^ |
| 7/17/2025 | Both | Amendment | Biosystem product Zinc acid (Zinc, sulfur, and organic acids) (4-6-4) | 9.35 | L ha^-1^ |
| 7/17/2025 | Both | Fertilizer | Nitrogen | 254 | kg ha^-1^ |

**Table S2** Soil properties measured in the free drainage (FD) and controlled drainage (CD) sections of the study field at 0-5 cm and 5-20 cm depths in fall and spring of 2020 and 2021, including organic matter, cation exchange capacity, pH, estimated nitrogen release, nitrate-nitrogen (NO_3_⁻–N), ammonium-nitrogen (NH_4_⁺-N), phosphorus (P), Bray II phosphorus, iron (Fe), aluminum (Al), sulfur (S), calcium (Ca), magnesium (Mg), potassium (K), sodium (Na), base saturation (Ca, Mg, K, Na, and other bases), hydrogen (H) saturation, boron (B), manganese (Mn), copper (Cu), zinc (Zn), phosphorus saturation ratio (PSR), and soil phosphorus storage capacity (SPSC).

|  | **Field and Depth** | | | |
| --- | --- | --- | --- | --- |
| **Parameter** | **Free Drainage (FD)** | | **Controlled Drainage (CD)** | |
|  | 0-5cm | 5-20cm | 0-5cm | 5-20cm |
| **Organic Matter (%)** | 2.5 | 2.2 | 3.1 | 2.8 |
| **Total Exchange Capacity (meq/100 g)** | 14 | 13 | 12 | 13 |
| **pH** | 7.7 | 7.6 | 7.6 | 7.4 |
| **Estimated Nitrogen Release (# N/acre)** | 70 | 63 | 80 | 75 |
| **NO_3_^-^-N (ppm)** | 8 | 12 | 7 | 13 |
| **NH_4_^+^-N (ppm)** | 3 | 3 | 2 | 3 |
| **P (mg/kg)** | 140 | 95 | 133 | 90 |
| **Bray II P (mg/kg)** | 152 | 105 | 157 | 109 |
| **Fe (mg/kg)** | 358 | 328 | 319 | 295 |
| **Al (mg/kg)** | 408 | 433 | 504 | 570 |
| **S (ppm)** | 10 | 10 | 10 | 10 |
| **Ca (mg/kg)** | 1949 | 1992 | 1619 | 1738 |
| **Mg (mg/kg)** | 296 | 284 | 337 | 356 |
| **K (mg/kg)** | 261 | 155 | 261 | 164 |
| **Na (mg/kg)** | 35 | 36 | 37 | 38 |
| **Ca (%)** | 70 | 72 | 66 | 68 |
| **Mg (%)** | 20 | 20 | 23 | 23 |
| **K (%)** | 5 | 3 | 6 | 3 |
| **Na (%)** | 1 | 1 | 1 | 1 |
| **Other Bases (%)** | 4 | 4 | 4 | 4 |
| **H(%)** | 0 | 0 | 0 | 0 |
| **B (mg/kg)** | 1 | 1 | 1 | 1 |
| **Mn (mg/kg)** | 24 | 22 | 20 | 15 |
| **Cu (mg/kg)** | 7 | 4 | 7 | 4 |
| **Zn (mg/kg)** | 7 | 5 | 8 | 6 |
| **PSR** | 0.22 | 0.14 | 0.18 | 0.11 |
| **SPSC** | -67 | -20 | -50 | 0 |

*****We sampled soils during both fall and spring in 2020 and 2021 according to methods adapted from Christopher et al., 2021 (Christopher et al., 2021). Briefly, we took soil cores at 5 m intervals along twelve 30 m transects that were evenly spaced throughout each field. One core per transect was taken using a bulk density slide hammer (AMS, American Falls, ID), while the remaining cores were taken with an Oakfield probe (Model H; Forestry Suppliers, Jackson, MS). For both types of cores, we subdivided each core into two depths (0-5 cm and 5-20 cm) then pooled and homogenized all cores within each transect. We stored samples in polyethylene bags and transported them to the laboratory on ice. We air dried, sieved (2 mm), and ground samples and then sent them to a commercial laboratory for analysis (Brookside Laboratories Inc; https://www.blinc.com/soil-testing-methods).

**Table S3** Percentage of time the controlled drainage (CD) system was managed during each water year (WY; October 1–September 30). Managed conditions were defined as the top of the DWM structure being set 100 cm below the field apex.

| **Water Year** | **Time Controlled Drainage was Managed (CD) (%)** |
| --- | --- |
| 2018 | 67.0 |
| 2019 | 64.5 |
| 2020 | 67.2 |
| 2021 | 99.8 |
| 2022 | 74.8 |
| 2023 | 50.5 |
| 2024 | 63.4 |
| 2025 | 76.8 |

**Table S4** Assay sequences and cycling conditions for targets. Target include *tetW* (assay developed by Knapp et al. 2009), *tetM* (assay developed by Peak et al. 2007), *ermB* (assay developed by Böckelmann et al. 2009), *sul1* (assay developed by McKinney et al. 2018). For each assay, forward and reverse primer sequences, probe sequences, amplicon length (bp), and dPCR cycling conditions are provided.

| **Target** | **Primers sequence (Forward and Reverse)** | **Probe sequence** | **Amplicon Length (bp)** | **Cycling conditions** |
| --- | --- | --- | --- | --- |
| *tetW* | F: GCAGAGCGTGGTTCAGTCT  R:GACACCGTCTGCTTGATGATAAT | HEX-TTCGGGATAAGCTCTCCGCCGA | 66 | All 95 ℃ for 2 min  40 cycles of 95 ℃ for 15 s  and 55 ℃ for 30 s |
| *tetM* | F: GGT TTC TCT TGG ATA CTT AAA TCA ATC R  R: CCA ACC ATA YAA TCC TTG TTC RC | 5HEX-AT GCA GTT A/ZEN/T GGA RGG GAT ACG CTA TGG Y/3IABkFQ | 88 |  |
| *ermB* | F:GGATTCTACAAGCGTACCTTGGA  R:GCTGGCAGCTTAAGCAATTGCT | FAM-CACTAGGGTTGCTCTTGCACACTCAAGTC | 92 |  |
| *sul1* | F:CCGTTGGCCTTCCTGTAAAG  R: TTGCCGATCGCGTGAAGT | 6FAM-CAGCGAGCCTTGCGGCGG | 67 |  |

**Table S5** Detection frequencies of antibiotic resistance genes (ARGs) in tile drainage water by drainage-management scenario. The number and percentage of tile-drain observations exceeding the target-specific limit of detection (LOD) or detection threshold are reported for each ARG and scenario. S2 represents managed controlled drainage with flow, S3 represents both tile drains unmanaged (free draining), and S4 represents the net effect comparison across management conditions. LODs/thresholds are reported in copies µL⁻¹ for *ermB* and *tetW* and as positive partitions for *sul1* and *tetM*.

| Scenario | Tile drains | # over LOD/  Threshold | % over LOD/  Threshold | Target | LOD/  Threshold | Units |
| --- | --- | --- | --- | --- | --- | --- |
| S3 | 14 | 3 | 21.4 | *ermB* | 0.296 | copies µL^-1^ |
| S2 | 8 | 2 | 25.0 | *ermB* | 0.296 | copies µL^-1^ |
| S4 | 22 | 5 | 22.7 | *ermB* | 0.296 | copies µL^-1^ |
| S3 | 14 | 7 | 50.0 | *tetW* | 0.262 | copies µL^-1^ |
| S2 | 8 | 7 | 87.5 | *tetW* | 0.262 | copies µL^-1^ |
| S4 | 22 | 14 | 63.6 | *tetW* | 0.262 | copies µL^-1^ |
| S3 | 14 | 5 | 35.7 | *sul1* | 2 | Positive partitions |
| S2 | 8 | 5 | 62.5 | *sul1* | 2 | Positive partitions |
| S4 | 22 | 10 | 45.5 | *sul1* | 2 | Positive partitions |
| S3 | 14 | 5 | 35.7 | *tetM* | 2 | Positive partitions |
| S2 | 8 | 7 | 87.5 | *tetM* | 2 | Positive partitions |
| S4 | 22 | 12 | 54.5 | *tetM* | 2 | Positive partitions |

**Table S6** Summary of generalized linear mixed model (GLMM) results for all measured response variables across drainage management scenarios (S2: Managed, S3: Unmanaged, S4: Net Effect). For each response, the table reports the model structure, error distribution and link function, random effect(s), response units, dispersion parameter (and Tweedie power parameter where applicable), estimated mean ratio (back-transformed ratio of estimated marginal means, treatment relative to reference level, obtained via emmeans::pairs(..., type = "response")) with associated 95% confidence interval, marginal R² (variance explained by fixed effects alone) and conditional R² (variance explained by fixed and random effects combined; Nakagawa & Schielzeth 2013), and *p*-value. For models with a Drain × Target interaction (ARG Concentration) or Drain × Season / Drain × depth interaction (Flashiness, Peak Counts, Soil moisture), estimated mean ratios and CIs are reported separately for each Target gene, season, or depth level below the overall model row. Abbreviations: ARG = antibiotic resistance gene; *tetW*, *tetM*, *ermB*, *sul1* = target ARG assays; SRP = soluble reactive phosphorus; NO_3_^⁻^-N = nitrate-nitrogen; NH_4_^⁺^-N = ammonium-nitrogen; CFU = colony-forming units; R-B Index (FI) = Richards-Baker Flashiness Index; VWC = volumetric water content. "ARG Concentration (without 02/28/2024)" rows repeat the corresponding model excluding samples collected on 2/28/2026 as a sensitivity check. Dashes (-) indicate values not applicable to that model or not reported for that row.

| **Response** | **Solute**  **or**  **Target** | **Scenario** | **Model Structure** | **GLM**  **Family** | **Random Effect** | **Units** | **Tweedie power parameter** | **Dispersion parameter/estimate** | **Estimated mean ratio**  **(CD/FD)** | **95% CI** | **Marginal R^2^** | **Conditional R^2^** | ***p*-value** |
| --- | --- | --- | --- | --- | --- | --- | --- | --- | --- | --- | --- | --- | --- |
| Water drainage yield | - | S2: Managed | Response ~ Drain + (1\|Year) | Gamma  (log link) | Year | L s^-1^ | - | 1.14 | 0.35 | 0.31-0.39 | 0.26 | 0.27 | **p<0.0001** |
|  |  | S3: Unmanaged |  | Tweedie |  |  | 1.51 | 1.11 | 0.861 | 0.79-0.94 | 0.007 | 0.217 | **p<0.001** |
|  |  | S4: Net Effect |  | Tweedie |  |  | 1.56 | 0.967 | 0.442 | 0.41-0.47 | 0.124 | 0.177 | **p<0.0001** |
| ARG Concentration | - | S2: Managed | Response ~ Drain × Target + (1\|Date) | Tweedie | Date | Gene copies L^-1^ | 1.275 | 175 | - | - | 0.133 | 0.999 | - |
|  | *tetW* |  |  |  |  |  | - | - | 1.071 | 0.91-1.26 | - | - | p>0.05 |
|  | *tetM* |  |  |  |  |  | - | - | 0.459 | 0.38-0.56 | - | - | **p<0.0001** |
|  | *ermB* |  |  |  |  |  | - | - | 0.873 | 0.58-1.33 | - | - | p>0.05 |
|  | *sul1* |  |  |  |  |  | - | - | 0.315 | 0.24-0.42 | - | - | **p<0.0001** |
| ARG Concentration | - | S3: Unmanaged | Response ~ Drain × Target + (1\|Date) | Tweedie | Date | Gene copies L^-1^ | 1.408 | 105 | - | - | 0.196 | 0.96 | - |
|  | *tetW* |  |  |  |  |  | - | - | 1.032 | 0.45-2.39 | - | - | p>0.05 |
|  | *tetM* |  |  |  |  |  | - | - | 1.109 | 0.40-3.08 | - | - | p>0.05 |
|  | *ermB* |  |  |  |  |  | - | - | 1.46 | 0.16-13.2 | - | - | p>0.05 |
|  | *sul1* |  |  |  |  |  | - | - | 0.801 | 0.28-2.28 | - | - | p>0.05 |
| ARG Concentration | - | S4: Net Effect | Response ~ Drain × Target + (1\|Date) | Tweedie | Date | Gene copies L^-1^ | 1.407 | 99.6 | - | - | 0.109 | 0.994 | - |
|  | *tetW* |  |  |  |  |  | - | - | 1.099 | 0.81-21.5 | - | - | p>0.05 |
|  | *tetM* |  |  |  |  |  |  |  | 0.537 | 0.3-0.88 |  |  | **p<0.001** |
|  | *ermB* |  |  |  |  |  |  |  | 0.914 | 0.47-1.77 |  |  | p>0.05 |
|  | *sul1* |  |  |  |  |  |  |  | 0.415 | 0.27-0.65 |  |  | **p<0.001** |
| ARG Yield | - | S2: Managed | Response ~ Drain + Target + (1\|Date) | Tweedie | Date | Gene copies ha^-1^ d^-1^ | 1.634 | 441 | 0.229 | 0.16-0.33 | 0.138 | 0.995 | **p<0.0001** |
|  |  | S3: Unmanaged |  |  |  |  | 1.564 | 3.20E+03 | 0.992 | 0.49-0.16 | 0.097 | 0.989 | p>0.05 |
|  |  | S4: Net Effect |  |  |  |  | 1.605 | 9.90E+02 | 0.265 | 0.18-0.40 | 0.123 | 0.981 | **p<0.0001** |
| Nutrient Concentration | SRP | S2: Managed | Response ~ Drain + (1\|Year) | Gamma  (log link) | Year | mg L^-1^ | - | 0.414 | 1.07 | 0.81-1.41 | 0.002 | 0.399 | p>0.05 |
|  |  | S3: Unmanaged |  |  |  |  |  | 0.878 | 0.808 | 0.55-1.2 | 0.016 | 0.114 | p>0.05 |
|  |  | S4: Net Effect |  |  |  |  |  | 0.701 | 0.921 | 0.71-2.19 | 0.003 | 0.202 | p>0.05 |
| Nutrient Concentration | NO_3_^-^ -N | S2: Managed | Response ~ Drain + (1\|Year) | Gamma  (log link) | Year | mg L^-1^ | - | 0.116 | 0.942 | 0.81-1.09 | 0.003 | 0.685 | p>0.05 |
|  |  | S3: Unmanaged |  |  |  |  |  | 0.157 | 0.948 | 0.80-1.12 | 0.002 | 0.638 | p>0.05 |
|  |  | S4:  Net Effect |  |  |  |  |  | 0.177 | 0.94 | 0.83-1.07 | 0.002 | 0.616 | p>0.05 |
| Nutrient Concentration | NH_4_^+^-N | S2: Managed | Response ~ Drain + (1\|Year) | Gamma  (log link) | Year | mg L^-1^ |  | 1.53 | 0.637 | 0.35-1.16 | 0.027 | 0.507 | p>0.05 |
|  |  | S3: Unmanaged |  |  |  |  |  | 2.11 | 0.653 | 0.34-1.27 | 0.027 | 0.327 | p>0.05 |
|  |  | S4:  Net Effect |  |  |  |  |  | 1.9 | 0.707 | 0.45-1.1 | 0.016 | 0.435 | p>0.05 |
| Nutrient Yield | SRP | S2: Managed | Response ~ Drain + (1\|Year) | Gamma  (log link) | Year | kg ha^-1^ d^-1^ | - | 2.23 | 0.481 | 0.25-0.94 | 0.082 | 0.286 | **p<0.05** |
|  |  | S3: Unmanaged |  | Tweedie |  |  | 1.857 | 0.808 | 0.521 | 0.27-1.01 | 0.063 | 0.122 | **p<0.0001** |
|  |  | S4:  Net Effect |  | Tweedie |  |  | 1.76 | 0.592 | 0.495 | 0.32-0.78 | 0.057 | 0.257 | **p<0.01** |
| Nutrient Yield | NO_3_^-^ -N | S2: Managed | Response ~ Drain + (1\|Year) | Tweedie | Year | kg ha^-1^ d^-1^ | - | 1.45 | 0.461 | 0.32-0.66 | 0.202 | 0.225 | **p<0.0001** |
|  |  | S3: Unmanaged |  |  |  |  | 1.723 | 0.0916 | 0.821 | 0.05-1.3 | 0.006 | 0.548 | p>0.05 |
|  |  | S4:  Net Effect |  |  |  |  | 1.631 | 0.0703 | 0.461 | 0.32-0.66 | 0.081 | 0.317 | **p<0.0001** |
| Nutrient Yield | NH_4_^+^-N | S2: Managed | Response ~ Drain + (1\|Year) | Gamma  (log link) | Year | kg ha^-1^ d^-1^ | - | 2.38 | 0.344 | 0.17-0.70 | 0.094 | 0.604 | **p<0.01** |
|  |  | S3: Unmanaged |  | Tweedie |  |  | 1.901 | 1.49 | 0.596 | 0.27-1.34 | 0.026 | 0.437 | p>0.05 |
|  |  | S4:  Net Effect |  | Tweedie |  |  | 1.817 | 0.993 | 0.561 | 0.33-0.95 | 0.024 | 0.486 | **p<0.05** |
| E.coli Concentration | - | S2: Managed | Response ~ Drain + (1\|Year) | Gamma  (log link) | Year | CFU 100 mL^-1^ | - | 1.33 | 0.762 | 0.52-1.12 | 0.008 | 0.62 | p>0.05 |
|  |  | S3: Unmanaged |  |  |  |  | - | 1.79 | 0.691 | 0.45-1.07 | 0.015 | 0.562 | p>0.05 |
|  |  | S4:  Net Effect |  |  |  |  | - | 2.2 | 0.535 | 0.38-0.75 | 0.032 | 0.609 | **p<0.0001** |
| E.coli Yield | - | S2: Managed | Response ~ Drain + (1\|Year) | Tweedie | Year | CFU ha^-1^ d^-1^ | 1.72 | 530 | 0.195 | 0.07-0.55 | 0.114 | 0.727 | **p<0.01** |
|  |  | S3: Unmanaged | Response ~ Drain |  |  |  | 1.71 | 1.47E+03 | 0.691 | 0.45-1.07 | - | - | p>0.05 |
|  |  | S4:  Net Effect | Response ~ Drain + (1\|Year) |  |  |  | 1.72 | 997 | 0.3 | 0.15-0.60 | 0.068 | 0.564 | **p<0.001** |
| Flashiness | - | Flashiness | Response ~ Drain*Season + (1\|Water Year) | Gamma  (log link) | Water Year | R-B Index (FI) | - | - | - | - | 0.313 | 0.569 | - |
|  |  |  | fall |  |  |  |  |  | 0.947 | 0.74-1.21 | - | - | p>0.05 |
|  |  |  | spring |  |  |  |  |  | 1.255 | 1.01-1.56 |  |  | **p<0.05** |
|  |  |  | summer |  |  |  |  |  | 1.299 | 1.04-1.62 |  |  | **p<0.05** |
|  |  |  | winter |  |  |  |  |  | 1.299 | 1.04-1.62 |  |  | **p<0.05** |
| Peak Counts | - | Peak Counts | Response ~ Drain*Season + (1\|Water Year) | Poisson  (log link) | Water Year | # | - | - | - | - | 0.27 | 0.498 | - |
|  |  |  | fall |  |  |  |  |  | 0.829 | 0.53-1.31 | - | - | p>0.05 |
|  |  |  | spring |  |  |  |  |  | 0.767 | 0.52-1.13 |  |  | p>0.05 |
|  |  |  | summer |  |  |  |  |  | 0.971 | 0.60-1.57 |  |  | p>0.05 |
|  |  |  | winter |  |  |  |  |  | 0.391 | 0.25-0.61 |  |  | **p<0.0001** |
| Soil moisture | - | 2021 |  | Gamma  (log link) | Month | VWV |  | 0.113 | - | - | 0.62 | 0.74 | - |
|  |  |  | 5 |  |  |  |  |  | 0.448 | 0.34–0.59 |  |  | **p<0.0001** |
|  |  |  | 10 |  |  |  |  |  | 0.426 | 0.32–0.56 |  |  | **p<0.0001** |
|  |  |  | 20 |  |  |  |  |  | 0.760 | 0.57–1.01 |  |  | p>0.05 |
|  |  |  | 30 |  |  |  |  |  | 1.307 | 0.99–1.73 |  |  | p>0.05 |
|  |  |  | 40 |  |  |  |  |  | 2.315 | 1.75–3.07 |  |  | **p<0.0001** |
|  |  |  | 50 |  |  |  |  |  | 1.170 | 0.88–1.55 |  |  | p>0.05 |
|  |  |  | 60 |  |  |  |  |  | 1.783 | 1.35–2.36 |  |  | **p<0.001** |
|  |  |  | 75 |  |  |  |  |  | 2.169 | 1.64–2.87 |  |  | **p<0.0001** |
|  |  |  | 100 |  |  |  |  |  | 0.908 | 0.69–1.20 |  |  | p>0.05 |
| Soil moisture | - | 2022 |  | Gamma  (log link) | Month | VWV |  | 0.042 | - | - | 0.536 | 0.851 | - |
|  |  |  | 5 |  |  |  |  |  | 0.510 | 0.43–0.60 |  |  | **p<0.0001** |
|  |  |  | 10 |  |  |  |  |  | 0.582 | 0.49–0.69 |  |  | **p<0.0001** |
|  |  |  | 20 |  |  |  |  |  | 0.811 | 0.69–0.96 |  |  | p>0.05 |
|  |  |  | 30 |  |  |  |  |  | 1.140 | 0.97–1.34 |  |  | p>0.05 |
|  |  |  | 40 |  |  |  |  |  | 1.336 | 1.13–1.57 |  |  | **p<0.0001** |
|  |  |  | 50 |  |  |  |  |  | 0.969 | 0.82–1.14 |  |  | p>0.05 |
|  |  |  | 60 |  |  |  |  |  | 1.556 | 1.32–1.83 |  |  | **p<0.001** |
|  |  |  | 75 |  |  |  |  |  | 2.142 | 1.82–2.53 |  |  | **p<0.0001** |
|  |  |  | 100 |  |  |  |  |  | 1.041 | 0.88–1.23 |  |  | p>0.05 |
| Soil moisture | - | 2023 | Response~Drain*  depth+ (1\|Month) | Gamma  (log link) | Month | VWV | - | 0.036 | - | - | 0.6 | 0.87 | - |
|  |  |  | 5 |  |  |  | - | - | 0.455 | 0.39–0.54 |  |  | **p<0.0001** |
|  |  |  | 10 |  |  |  | - | - | 0.589 | 0.50–0.70 |  |  | **p<0.0001** |
|  |  |  | 20 |  |  |  | - | - | 0.731 | 0.62–0.86 |  |  | **p<0.0001** |
|  |  |  | 30 |  |  |  | - | - | 1.052 | 0.89–1.24 |  |  | p>0.05 |
|  |  |  | 40 |  |  |  | - | - | 0.841 | 0.71–0.99 |  |  | **p<0.05** |
|  |  |  | 50 |  |  |  | - | - | 0.867 | 0.73–1.02 |  |  | p>0.05 |
|  |  |  | 60 |  |  |  | - | - | 1.290 | 1.09–1.52 |  |  | **p<0.01** |
|  |  |  | 75 |  |  |  | - | - | 1.712 | 1.45–2.02 |  |  | **p<0.0001** |
|  |  |  | 100 |  |  |  | - | - | 1.091 | 0.92–1.29 |  |  | p>0.05 |
| ARG Concentration  (without 02/28/2024) | - | S2: Managed | Response ~ Drain × Target + (1\|Date) | Tweedie | Date | Gene copies L^-1^ | 1.24 | 129 |  |  | 0.75 | 0.918 |  |
|  | *tetW* |  |  |  |  |  |  | - | 3.268 | 1.54-6.95 |  |  | p<0.05 |
|  | *tetM* |  |  |  |  |  |  | - | 1.464 | 0.75-2.88 |  |  | p>0.05 |
|  | *ermB* |  |  |  |  |  |  | - | 0.981 | 0.10-10.2 |  |  | p>0.05 |
|  | *sul1* |  |  |  |  |  |  | - | 1.678 | 0.68-4.12 |  |  | p>0.05 |
| ARG Concentration (without 02/28/2024) | - | S3: Unmanaged | Response ~ Drain × Target + (1\|Date) | Tweedie | Date | Gene copies L^-1^ | 1.41 | 106 | - | - | 0.196 | 0.96 | p>0.05 |
|  | *tetW* |  |  |  |  |  |  | - | 1.032 | 0.45-2.39 | - | - | p>0.05 |
|  | *tetM* |  |  |  |  |  |  | - | 1.109 | 0.40-3.08 | - | - | p>0.05 |
|  | *ermB* |  |  |  |  |  |  | - | 1.46 | 0.16-13.2 | - | - | p>0.05 |
|  | *sul1* |  |  |  |  |  |  | - | 0.801 | 0.28-2.28 | - | - | p>0.05 |
| ARG Concentration (without 02/28/2024) | - | S4: Net Effect | Response ~ Drain × Target + (1\|Date) | Tweedie | Date | Gene copies L^-1^ | 1.2 | 92.1 |  |  | 0.259 | 0.949 | - |
|  | *tetW* |  |  |  |  |  |  |  | 1.41 | 0.75-2.67 | - | - | p>0.05 |
|  | *tetM* |  |  |  |  |  |  |  | 1.26 | 0.62-2.59 | - | - | p>0.05 |
|  | *ermB* |  |  |  |  |  |  |  | 1.37 | 0.26-7.37 | - | - | p>0.05 |
|  | *sul1* |  |  |  |  |  |  |  | 1.01 | 0.46-2.22 | - | - | p>0.05 |
| ARG Yield (without 02/28/2024) | - | S2: Managed | Response ~ Drain + Target + (1\|Date) | Tweedie | Date | Gene copies ha^-1^ d^-1^ | 1.35 | 1.77e+04 | 0.57 | 0.42-0.79 | 0.372 | 0.977 | **p<0.001** |
|  |  | S3: Unmanaged |  |  |  |  | 1.56 | 3.2e+03 | 0.88 | 0.49-1.59 | 0.097 | 0.989 | p>0.05 |
|  |  | S4: Net Effect |  |  |  |  | 1.57 | 2.14e+03 | 0.762 | 0.50-1.17 | 0.125 | 0.983 | p>0.05 |

**Table S7** Cumulative water yield (mm) and percent difference in yields by water year, and cumulative nutrient (NH_4_^+^-N: ammonium-nitrogen; NO_3_^-^-N: nitrate-nitrogen; SRP: soluble reactive phosphorus) yield. Values by water year for controlled drainage (CD) and free drainage (FD) fields.

|  |  |  |  | **NO_3_^-^-N Yield (kg)** | | **NH_4_^+^-N (kg)** | | **SRP (kg)** | |
| --- | --- | --- | --- | --- | --- | --- | --- | --- | --- |
| **Water Year** | **Free drainage (FD) yield (mm)** | **Controlled drainage (CD) yield (mm)** | **Percentage Difference** | **Free Drainage (FD)** | **Controlled Drainage (CD)** | **Free Drainage (FD)** | **Controlled Drainage (CD)** | **Free Drainage (FD)** | **Controlled Drainage (CD)** |
| 2018 | 515 | 266 | 48 | - | - | - | - | - | - |
| 2019 | 636 | 313 | 51 | 99.2 | 41.9 | 0.14 | 0.09 | 0.63 | 0.31 |
| 2020 | 687 | 441 | 36 | 74.4 | 39.9 | 0.21 | 0.32 | 0.56 | 0.5 |
| 2021 | 291 | 60 | 79 | 111.8 | 19.6 | 0.01 | 0 | 0.2 | 0.09 |
| 2022 | 635 | 247 | 61 | 120.1 | 57.7 | 0.09 | 0.07 | 0.32 | 0.13 |
| 2023 | 615 | 354 | 42 | 98.3 | 52.1 | 0.12 | 0.07 | 0.27 | 0.16 |
| 2024 | 651 | 311 | 52 | 118.4 | 45.7 | 0.39 | 0.09 | 0.76 | 0.31 |
| 2025 | 592 | 106 | 82 | 39.9 | 5.4 | 0.05 | 0.01 | 0.3 | 0.05 |

**Table S8** Cumulative water and nutrient yields by water year. Cumulative water yield (mm) and percent difference in water yield between controlled drainage (CD) and free drainage (FD) are reported for each water year. Cumulative nutrient yields are reported for ammonium-N (NH_4_^+^-N), nitrate-N (NO_3_^-^-N), and soluble reactive phosphorus (SRP) for CD and FD fields.

| **Response**  **variable** | **Metric** | **Unit** | **Managed but flowing (S2)** | | | **Unmanaged (S3)** | | | **Net Effect (S4)** | |
| --- | --- | --- | --- | --- | --- | --- | --- | --- | --- | --- |
|  |  |  | **FD** | **CD** | **FD** | | **CD** | **FD** | | **CD** |
| Water | Daily drainage yield | mm | 3.43 ± 0.20 | 1.21 ± 0.08 | 1.94 ± 0.28 | | 1.67 ± 0.24 | 1.69 ± 0.16 | | 0.74 ± 0.07 |
| NO_3_^-^-N | Concentration | mg L^-1^ | 17.2 ± 3.17 | 16.2 ± 2.98 | 11.5 ± 2.49 | | 10.9 ± 2.36 | 15.3 ± 2.87 | | 14.3 ± 2.70 |
| NO_3_^-^-N | Daily yield | kg ha^-1^ | 1.87x10⁻⁴ ± 4.17x10⁻⁵ | 7.17x10⁻⁵ ± 1.46x10⁻⁵ | 6.59x10⁻⁵ ± 3.18x10⁻⁵ | | 5.41x10⁻⁵ ± 2.62x10⁻⁵ | 6.67x10⁻⁵ ± 1.79x10⁻⁵ | | 3.08x10⁻⁵ ± 8.60x10⁻⁶ |
| NH_4_^+^-N | Concentration | ug L^-1^ | 28.7 ± 11.8 | 18.3 ± 7.43 | 28.5 ± 11.1 | | 18.6 ± 7.32 | 24.3 ± 8.82 | | 17.2 ± 6.17 |
| NH_4_^+^-N | Daily yield | kg ha^-1^ | 2.56x10⁻⁴ ± 1.38x10⁻⁴ | 8.83x10⁻⁵ ± 4.64x10⁻⁵ | 2.26x10⁻⁴ ± 1.33x10⁻⁴ | | 1.35x10⁻⁴ ± 7.88x10⁻⁵ | 8.66x10⁻⁵ ± 4.24x10⁻⁵ | | 4.86x10⁻⁵ ± 2.41x10⁻⁵ |
| SRP | Concentration | µg L^-1^ | 67.0 ± 10.9 | 61.7 ± 10.1 | 63.1 ± 11.6 | | 51.0 ± 9.34 | 73.0 ± 14.7 | | 77.9 ± 15.9 |
| SRP | Daily yield | kg ha^-1^ | 9.67x10⁻⁴ ± 3.24x10⁻⁴ | 4.65x10⁻⁴ ± 1.53x10⁻⁴ | 6.33x10⁻⁴ ± 2.91x10⁻⁴ | | 3.30x10⁻⁴ ± 1.49x10⁻⁴ | 3.81x10⁻⁴ ± 1.08x10⁻⁴ | | 1.89x10⁻⁴ ± 5.54x10⁻⁵ |
| *E. coli* | Concentration | CFU 100 mL^-1^ | 22.4 ± 9.56 | 17.1 ± 7.27 | 33.1 ± 14.8 | | 22.9 ± 10.3 | 45.9 ± 21.0 | | 24.6 ± 11.4 |
| *E. coli* | Daily yield | CFU ha^-1^ | 1.09x10⁷ ± 9.06x10⁶ | 2.13x10⁶ ± 1.79x10⁶ | 2.08x10⁷ ± 7.27x10⁶ | | 1.41x10⁷ ± 5.22x10⁶ | 9.11x10⁶ ± 5.67x10⁶ | | 2.73x10⁶ ± 1.77x10⁶ |
| *tetW* | Concentration | Gene copies L^-1^ | 1,224 ± 1,290 | 1,264 ± 1,330 | 29,025 ± 39,900 | | 31,094 ± 42,800 | 3,732 ± 3,560 | | 4,101 ± 3,910 |
| *tetM* | Concentration | Gene copies L^-1^ | 590 ± 636 | 655 ± 704 | 27,206 ± 37,400 | | 12,490 ± 17,200 | 3,304 ± 3,150 | | 1,773 ± 1,690 |
| *ermB* | Concentration | Gene copies L^-1^ | 38.5 ± 50.7 | 56.2 ± 70.9 | 2,289 ± 3,170 | | 1,999 ± 2,770 | 269 ± 263 | | 246 ± 241 |
| *sul1* | Concentration | Gene copies L^-1^ | 661 ± 707 | 530 ± 571 | 15,014 ± 20,700 | | 4,737 ± 6,540 | 1,953 ± 1,860 | | 811 ± 780 |
| *tetW* | Daily yield | Gene copies ha^-1^ | 5.60x10⁸ ± 9.17x10⁸ | 1.28x10⁸ ± 2.10x10⁸ | 1.66x10⁷ ± 2.49x10⁷ | | 1.47x10⁷ ± 2.20x10⁷ | 7.20x10⁷ ± 6.35x10⁷ | | 1.90x10⁷ ± 1.69x10⁷ |
| *tetM* | Daily yield | Gene copies ha^-1^ | 4.55x10⁸ ± 7.44x10⁸ | 1.04x10⁸ ± 1.70x10⁸ | 8.83x10⁶ ± 1.34x10⁷ | | 7.79x10⁶ ± 1.18x10⁷ | 5.18x10⁷ ± 4.58x10⁷ | | 1.37x10⁷ ± 1.22x10⁷ |
| *ermB* | Daily yield | Gene copies ha^-1^ | 3.92x10⁷ ± 6.47x10⁷ | 8.97x10⁶ ± 1.48x10⁷ | 7.35x10⁵ ± 1.15x10⁶ | | 6.49x10⁵ ± 1.02x10⁶ | 4.63x10⁶ ± 4.22x10⁶ | | 1.22x10⁶ ± 1.12x10⁶ |
| *sul1* | Daily yield | Gene copies ha^-1^ | 2.38x10⁸ ± 3.89x10⁸ | 5.43x10⁷ ± 8.91x10⁷ | 9.49x10⁶ ± 1.43x10⁷ | | 8.37x10⁶ ± 1.26x10⁷ | 3.42x10⁷ ± 3.02x10⁷ | | 9.06x10⁶ ± 8.07x10⁶ |
